# Characterizing Cell-Free Transcription and Translation Dynamics with Nucleic Acid-Based Assays

**DOI:** 10.1101/2025.09.10.675373

**Authors:** Fernanda Piorino, Chad A. Sundberg, Elizabeth A. Strychalski, Eugenia Romantseva

## Abstract

Characterization of cell-free expression (CFE) systems must expand beyond single spectrophotometric measurements of a green fluorescent protein to provide meaningful metrics of system performance during a CFE reaction and enable the development of predictable and reproducible CFE technologies. To date, comprehensive characterization of these systems has posed a formidable measurement challenge, as it requires time-course measurements of reactions involving endogenous components in addition to transcription and translation of a target genetic circuit added exogenously to the CFE reaction. To provide more informative characterization that is still easy to conduct and complements current practices, we demonstrate a measurement framework for transcription and translation dynamics. We use different nucleic acid templates to characterize a suite of *Escherichia coli* extracts prepared in house, as well as extracts and reconstituted systems available commercially. Notably, we include measurements of low-performing systems to assess the sensitivity of our measurement framework and elucidate metrics indicative of system performance. For all these CFE systems, we compute reaction metrics to enable quantitative comparison. We believe this is an accessible measurement framework that can complement existing characterization, provide informative data for developing CFE technologies, and be adopted for routine characterization.

## Introduction

Realizing the full potential of cell-free expression (CFE) systems(*1–3*) requires measurement tools and methods to share fit-for-purpose metrics, enable effective collaboration, build on successes in the field, and improve confidence in CFE technologies(*4, 5*). Such a measurement framework for CFE systems would provide the foundation for precision engineering and implementation of these systems beyond the bench. However, the field still needs to identify and agree upon metrics indicative of CFE performance and suitable tools and assays to make these measurements. Whether based on crude lysates or purified proteins, CFE systems have many residual components from the host organism, which, along with exogenous reagents, participate in reactions even beyond the transcription and translation of a target genetic circuit. As these components interact, the composition and dynamics of the system change substantially over the course of a CFE reaction. Because of this complexity, characterizing CFE systems remains an unmet measurement challenge.

Current characterization practices fall short of measuring the complexity of CFE systems. A common preliminary assessment of crude lysate quality involves measuring the total concentration of endogenous proteins in the lysate, for example, via a Bradford Assay(*6*), but does not determine the identity or activity of proteins that may affect CFE. To verify expression of a target or model protein during a CFE reaction, characterization efforts typically include gel electrophoresis(*7–10*), which assesses protein expression visually and can provide quantitative information on band intensity when coupled with suitable software(*11, 12*). A widespread metric of reaction productivity is the maximum or endpoint fluorescence of a green fluorescent protein (GFP) generated during a CFE reaction(*5*). Because published studies typically report measurements in arbitrary units and use DNA templates with different components and GFP variants, we often cannot compare fluorescence measurements across or even within laboratories. In addition, GFP fluorescence is an uncertain predictor of the expression of proteins more structurally and functionally complex than common GFP variants, which are genetically engineered to mature rapidly, fold efficiently, and remain photostable. Although insufficient for full characterization of CFE systems, measurements of GFP could contribute to a larger measurement framework that considers the benefits and limitations of these measurements. Anecdotally, several laboratories in academia and industry have reported internal quality control strategies, although these tend to be application specific and considered intellectual property. To date, no quality control assays, other than gel electrophoresis and measurements of total protein levels and maximum GFP fluorescence, have seen wide adoption by users of CFE systems or could easily integrate into current CFE workflows.

Characterizing CFE systems requires measurements capturing two classes of reactions. The first and most frequent focus of quality control efforts focuses on the transcription and translation of a target genetic circuit encoded in a nucleic acid template added to the CFE reaction. The second consists of background reactions beyond transcription and translation of a target genetic circuit and is driven by residual components, such as small molecules and proteins, and pathways that are native to the host organism and remain in the system after cell lysis(*13–15*). Measuring this endogenous metabolism is resource- and time-intensive and thus unsuitable for routine CFE characterization. Therefore, we propose a measurement framework with an initial focus on measurements of transcription and translation dynamics over the course of a CFE reaction.

In our measurement framework, we leverage nucleic acid templates to measure transcription and translation dynamics. With nucleic acid templates, our characterization assays remain complementary to current measurements with GFP. However, our assays surpass a single fluorescence measurement by incorporating time-course measurements of both protein and RNA. Using different genetic parts and types of nucleic acids, we measure and decouple transcription and translation over the course of CFE reactions. In this way, we derive quantitative metrics to assess system dynamics and performance. We anticipate this characterization approach to be facile to implement, as users of CFE systems have prior experience preparing and handling nucleic acid templates for CFE. In addition, nucleic acid sequences can be shared easily to help align the protocols and genetic parts used for CFE measurements.

Our proposed metrology framework provides routine benchmarking of CFE to complement measurements of GFP fluorescence and enable comparison of different systems. We include time-course measurements of both a GFP variant and an RNA reporter to characterize CFE systems from different host strains, lysate preparation conditions, reaction formulations, vessels, and volumes. We provide assay results for both reconstituted and lysate-based *Escherichia coli* (*E. coli*) CFE systems used widely by the community. Importantly, we include low-performing systems to help elucidate metrics indicative of system performance and determine whether our measurement framework captures suboptimal performance. Finally, we assess the utility of nucleic acids as measurement tools and discuss the state of measurement assurance for CFE, identifying additional gaps in current characterization and opportunities for future efforts. We believe this work is valuable to identifying key metrics, guiding protocol documentation, scoping future characterization efforts, and generally improving our understanding of CFE systems, as well as their accessibility and reproducibility.

## Results

### Measurement tools and measurands

We used different nucleic acid templates to decouple measurements of transcription and translation dynamics (**Figure 1A**). We selected plasmid pJL1 (Addgene #69496)(*16*) as the backbone for these templates, due its prevalence in published CFE studies. This plasmid expresses the protein reporter of our assays, superfolder GFP (sfGFP), using the consensus T7 promoter and terminator for transcription and T7 phage’s gene 10 ribosomal binding site (RBS) for translation(*17*). To measure transcription dynamics, we modified pJL1 to generate two additional plasmids, pFP34 and pFP35, by transcriptionally fusing sfGFP with a fluorogenic Pepper RNA aptamer. Upon transcription, the Pepper aptamer binds to a dye molecule, HBC620, to allow red fluorescence(*18*). Initially engineered to visualize live cells(*18*), Pepper aptamers have also been harnessed in RNA-based sensors(*19, 20*). Compared to other commonly used aptamers, Pepper aptamers are more photostable and less sensitive to magnesium ions, do not require potassium ions for folding, and use a dye safer to the user(*18, 21*). Each of pFP34 and pFP35 contains a Pepper dimer grafted onto an RNA scaffold that improves the fluorescence signal(*21*). In pFP34, the Pepper aptamer includes both a tRNA and a F30 scaffold (tDF30ppr); in plasmid pFP35, it includes only the F30 scaffold (DF30ppr), with ‘D’ denoting a Pepper dimer (**Figure 1B**). Upon binding to HBC620, tDF30ppr exhibits greater *in vivo* fluorescence than DF30ppr, likely due to enhanced stability, although the exact reason is unclear(*21*). Because the two scaffolds differ in signal, we included both here, expecting DF30ppr to be more sensitive to differences across CFE systems and tDF30ppr to enable reliable measurements even in low-performing systems. pJL1, pFP34, and pFP35 provide the basis for all other nucleic acid templates used in this manuscript; each figure specifies any changes to these templates, and **Supplementary File 2** includes annotated sequences.

**Figure 1.**
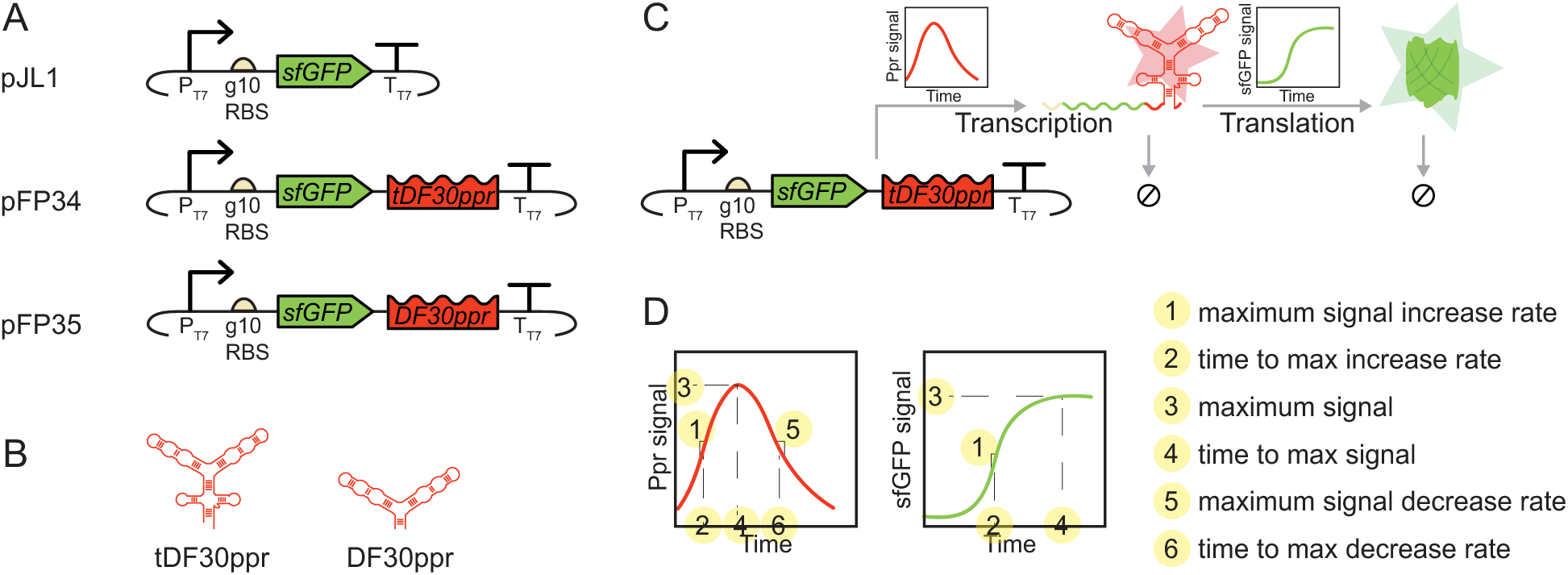
Tools and measurements to demonstrate our measurement framework. **(A)** Nucleic acid templates (pJL1, pFP34, and pFP35) used for time-course measurements of transcription and translation, with sfGFP as the protein reporter and the Pepper RNA aptamer as the RNA reporter. Based on pJL1, these plasmids use the T7 promoter and terminator for transcription, and T7 phage’s gene 10 (g10) ribosomal binding site (RBS) for translation. pFP34 and pFP35 have different Pepper aptamer scaffolds: pFP34 contains tDF30ppr, with both tRNA and F30 scaffolds, and pFP35 contains DF30ppr, with the F30 scaffold only. **(B)** Predicted secondary structures of the Pepper RNA aptamer in the two different RNA scaffolds, adapted from Mumbleau *et al*. (*21*). **(C)** Representative time-course RNA and protein measurements. With pFP34 as the template, for example, we monitor an increase in red fluorescence as tDF30ppr is transcribed, folds, and binds its cognate dye (HBC620), and a decrease in red fluorescence as degradation of tDF30ppr (indicated by ‘⦸’) exceeds transcription, folding, and binding. Similarly, we monitor an increase in green fluorescence as sfGFP is translated, although a decrease in green fluorescence is not expected in a protease-deficient host organism like *E. coli* BL21. **(D)** CFE metrics derived from time-course transcription and translation measurements. The values of these metrics are expected to change across CFE systems, nucleic acid templates, vessels, and scales. **Supplementary File 1** describes each metric and their calculation in more detail, and **Supplementary File 2** includes metrics values for all experiments.

These templates allowed simultaneous measurements of transcription and translation dynamics over the course of a CFE reaction, as Pepper mRNA and sfGFP were generated and degraded (**Figure 1C**). We derived different metrics for quantitative characterization of transcription and translation (**Figure 1D**). In crude lysates, which contain ribonucleases, mRNA dynamics first exhibit an increase in signal—resulting from transcription, folding, and binding of the Pepper aptamer to its cognate dye—followed by a decrease in signal as mRNAs degrade. From Pepper measurements, we thus computed the maximum signal increase rate, maximum signal, maximum signal decrease rate, and their respective time points. From sfGFP measurements, we derived the same metrics except for those associated with signal decrease, as sfGFP is highly stable(*22, 23*) and protease activity is generally low in lysates derived from the protease-deficient BL21 strain(*24*). To enable data comparability across laboratories, we reported fluorescence data in Molecules of Equivalent Soluble Fluorochrome (MESF) based on calibration curves for two commercially available fluorochromes: the green NIST-traceable fluorescein standard (NFS) as a metric of sfGFP signal and the red Atto 590 dye as a metric of HBC620-bound Pepper mRNA signal (**Figure S1**). To decouple transcription and translation, we further modified pFP34 and pFP35. For example, we removed the RBS to focus on transcription only and used mRNAs instead of DNA templates to bypass transcription.

To demonstrate the usability of our measurement framework, we tested host strains, lysate preparation conditions, reagents, and reaction formulations relevant to applications from basic scientific research to biomanufacturing. We selected *E. coli* as our host organism, because it is the most established for CFE, with widely reported protocols(*6, 25, 26*), data, and commercially available systems, in both lysate and reconstituted formats. To study lysate-based systems, which include a supernatant from lysed cells and preserve some endogenous material, we tested lysates prepared in house (**Table 1**) and available commercially (NEBExpress, New England Biolabs). Here, we refer to all lysates tested as ‘extracts’ to account for post-lysis processing. To study reconstituted CFE systems, which consist of a defined mixture of purified proteins that support transcription and translation, we tested the commercially available PURExpress (New England Biolabs), based on PURE(*27, 28*). Most of our in-house extracts derive from BL21 Star (DE3), a strain suitable for high-yield protein expression, because it is protease deficient and includes an IPTG-inducible T7 RNA polymerase (RNAP) expression cassette(*24*). The Star mutation denotes a truncated RNase E lacking the domain required for RNA degradation, and thus confers the bacterium higher RNA stability(*29*). **Table 1** lists the main differences among the preparation conditions for the in-house extracts tested, including the formulation of the cell resuspension buffer (S30 buffer, an acetate-based buffer(*25*), versus S30A, a glutamate-based buffer(*30, 31*)), the harvest time, and whether the preparation includes a runoff reaction (*32, 33*). Despite differences in the strains and methods used to prepare extracts, the total endogenous protein content of all extracts did not differ by more than 15 % (**Figure S2**), so differences in performance could not be attributed exclusively to the availability of endogenous machinery, such as proteins that enable transcription and translation. Unless specified, we ran reactions in batch mode, using the PANOx-SP(*31*) energy system at 10 µL volumes in a clear 384-well plate with an optically clear, flat bottom incubated at 37 °C in a multimode plate reader without shaking. We did not aim to explore different reaction formulations and formats rigorously, which have been studied elsewhere(*31, 34–37*) and could be the subject of future studies. Section III of **Supplementary File 1** provides more detail on the differences in genotypes, extract preparation conditions, and reaction formulations and formats among the systems tested.

**Table 1.** Characteristics of the *E. coli* extracts prepared in house used in this work, including the host organism, harvest time, cell resuspension buffer, and post-lysis processing. ‘+’ indicates that the extract preparation protocol included the reagent or step; ‘-’ indicates omission of the step or reagent.

| Extract | Strain | IPTG | Harvest | Buffer | Sonicator | Runoff | Relevant figure |
| --- | --- | --- | --- | --- | --- | --- | --- |
| EXT3 | BL21 Star (DE3) | + | Mid-exponential | S30 | Q125 | + | 2A, 3A, 4, 5, 6, 7, S3, S4B, S5, S8, S9, S11, S12, S13 |
| EXT4 | BL21 Star (DE3) | + | Mid-exponential | S30 | Q125 | + | S13 |
| EXT10 | BL21 Star (DE3) | + | Mid-exponential | S30 | Q700 (24-tip) | + | 4B, S9A |
| EXT12 | BL21 (DE3) | + | Mid-exponential | S30 | Q125 | + | 4D, 5B-C, S9C |
| EXT14 | BL21 Star (DE3) | + | Mid-exponential | S30A (pH 10) | Q125 | + | 4C, 5B-C, S9B |
| EXT15 | BL21 Star (DE3) | + | Late-exponential | S30 | Q125 | + | S10 |
| EXT16 | BL21 Star (DE3) | + | Late-exponential | S30 | Q125 | + | S10 |
| EXT17 | BL21 Star (DE3) | + | Mid-exponential | S30A | Q125 | + | 4B, 5B-C, S9A |
| EXT18 | BL21 Star (DE3) | + | Mid-exponential | S30A | Q125 | - | 4B, 5B-C, S9A |
| EXT19 | BL21 Star (DE3) | + | Stationary | S30 | Q125 | + | 4C, 5B-C, S9B |
| EXT20 | BL21 Star (DE3) | + | Mid-exponential | S30 | Q125 | - | 4B, 5B-C, S9A |
| EXT23 | BL21 Star (DE3) | - | Mid-exponential | S30 | Q125 | + | 4B, S9A |

### Baseline transcription and translation measurements

To establish baseline transcription and translation measurements as an internal reference for comparison with other systems, we first applied our measurement framework to assess CFE in EXT3, a BL21 Star (DE3) extract prepared under commonly reported conditions(*6, 25*). We selected a DNA concentration of 5 nmol/L for our assays to maximize sfGFP signal (**Figure S3**).

The transcriptional fusion of sfGFP with Pepper had weak effects on sfGFP expression, slightly speeding up expression and leading to small decrease in the maximum sfGFP signal from pFP34 relative to pJL1 (**Figure 2A, left**). Whereas sfGFP expression dynamics were similar from the two plasmids, Pepper mRNA dynamics were markedly different, with much higher signal from pFP34 than pFP35 during the reaction (**Figure 2A, center** vs. **Figure 2A, right**). With pFP34 and pFP35, the Pepper signal increased immediately, such that the time between the first two measurements was sufficient for the aptamer to transcribe, fold, bind HBC620, and generate a measurable signal. The two constructs reached a maximum rate of signal increase at the same time, although this rate was 17 % higher from pFP34 than pFP35. Pepper measurements from pFP34 also exhibited higher maximum and endpoint Pepper signals and a 2.2-fold lower maximum rate of signal decrease. Consistent with tDF30ppr having higher fluorescence *in vivo*(*21*), these results point to tDF30ppr being more stable than DF30ppr.

**Figure 2.**
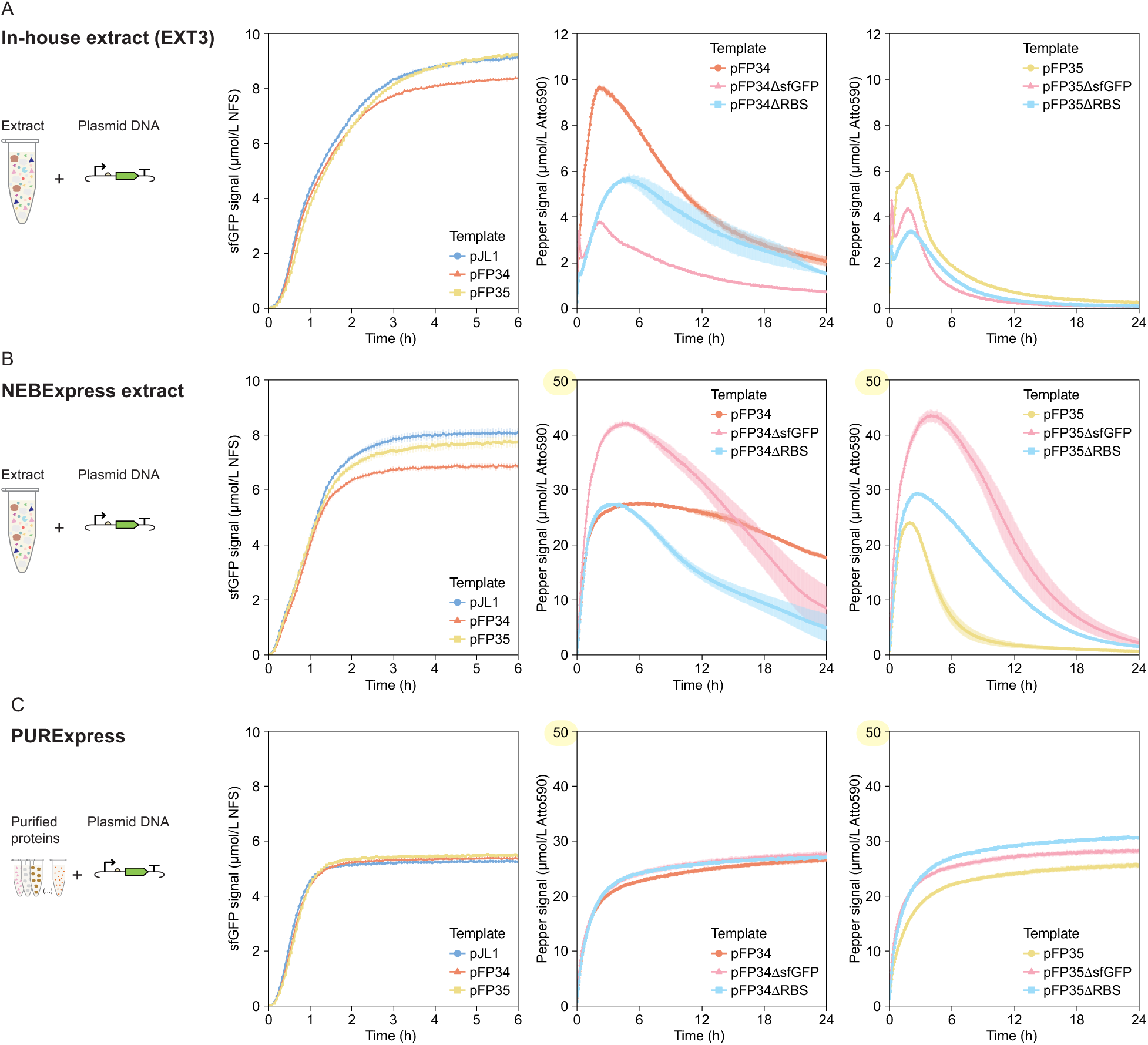
Baseline measurements of transcription and translation dynamics from DNA templates in **(A)** EXT3, **(B)** NEBExpress, and **(C)** PURExpress. All reactions include DNA templates at a concentration of 5 nmol/L. Note the different y-axis for Pepper measurements in the three CFE systems. sfGFP measurements are reported in Molecules of Equivalent Soluble Fluorochrome (MESF) of a NIST-traceable fluorescein standard (NFS) and are shown for only the first 6 h of the reaction, because the signal remains constant until 24 h. Pepper mRNA measurements are reported in MESF of Atto 590. The error bars indicate the standard deviation of three technical replicates.

To decouple measurements of transcription from translation, we removed either the RBS or the *sfGFP* gene from pFP34 and pFP35—generating pFP34ΔsfGFP, pFP35ΔsfGFP, pFP34ΔRBS, and pFP35ΔRBS—, modifications that affected transcription dynamics adversely. Removing *sfGFP* from pFP34 reduced the maximum Pepper signal increase rate 5.5-fold and the maximum signal 2.5-fold but caused only a 4 min lag to reach the peak signal (**Figure 2A, center**). While removing the RBS from pFP34 was not as detrimental to transcription as removing *sfGFP*, this modification delayed the maximum Pepper signal by almost 3 h relative to pFP34 (**Figure 2A, center**). The opposite trend was observed for the DF30ppr-encoding templates: pFP35ΔsfGFP reached a maximum signal higher than that of pFP35ΔRBS, and the reduction in maximum signal relative to pFP35 was not as pronounced (**Figure 2A, right**). Again, measurements with pFP35ΔRBS reached a maximum signal at a later time than with pFP35 and pFP35ΔsfGFP, although with a shorter lag than observed for the tDF30ppr-encoding templates. As expected, the modified templates did not generate sfGFP above background levels, because they lacked an RBS or the *sfGFP* gene (data not shown).

Although removing *sfGFP* or the RBS from nucleic acid templates did not enable full decoupling of transcription and translation, these modifications highlighted the importance of accounting for the genetic context when using nucleic acids to characterize CFE. Because transcription and translation share resources, we anticipated bypassing translation to enhance transcription. However, the measured effect on these processes depended on the components of the nucleic acid template and was subsequently masked by other processes active in the extract. In pFP34 and pFP35, *sfGFP* was upstream of Pepper and thus was transcribed first; we expected the ΔsfGFP template to exhibit faster Pepper signal increase because the lack of *sfGFP* would allow the aptamer to be transcribed immediately and the T7 RNAP to proceed to other DNA templates. However, it is possible that Pepper folded into a more stable conformation when transcriptionally fused with *sfGFP* or that the presence of *sfGFP* helped to protect the mRNA transcript from degradation by ribonucleases. The lack of translation in ΔRBS templates was also detrimental to Pepper signal, perhaps because the formation of a mRNA-ribosome complex protected the mRNA from degradation(*38*). For pFP34, removing the RBS affected transcriptional longevity; the Pepper signal from pFP34ΔRBS remained longer in a regime of signal increase. From a design standpoint, these results could inform deliberate modifications to nucleic acid templates to improve CFE for a given application. For example, including an RBS or a dummy gene in the DNA template could improve the RNA signal in some extracts; similarly, removing an extraneous RBS could improve RNA signal longevity.

To further characterize our measurement framework, we made the same measurements in commercially available CFE systems, in both lysate and reconstituted formats. An increasing number of stakeholders is interested in purchasing these systems to circumvent the higher variability and the time and resource investment associated with preparing CFE systems. Commercial reconstituted systems, most of which are based on the *E. coli*-derived PURE(*27, 28*), are particularly appealing, because they are difficult to prepare in house and, unlike lysates, have a defined composition that facilitates computational modeling(*39–42*) for engineering CFE systems. Whether reconstituted or lysate-based, commercially available CFE systems are often subject to trade secrets, so details on the host organism, preparation, and reaction formulation may be proprietary. Characterizing these systems, especially relative to those prepared in house, could inform which system to use for a given application, enable study of processes like RNA degradation that are more prominent in one type of system than another, and improve our general understanding of CFE systems.

We applied our measurement framework to characterize transcription and translation dynamics in NEBExpress, a commercially available *E. coli* extract. Compared with EXT3, translation in NEBExpress was slower and reached a lower maximum sfGFP signal, but was slightly more sensitive to the Pepper aptamer scaffold (**Figure 2B, left**). Transcription in NEBExpress was striking: the maximum Pepper signals and rates of signal increase were substantially higher in NEBExpress than in EXT3 from both pFP34 and pFP35 (**Figure 2B, center and right**). The Pepper signal from pFP34 was also stable: the signal decreased by only 33 % from its maximum value at 24 h. With pFP35, the Pepper signal reached a maximum value 5.0-fold higher and only 30 min later than in EXT3, so NEBExpress supported signal increase—whether by transcription, folding, or binding to HBC620—for longer (**Figure 2B, right**). After reaching a maximum, however, the Pepper signal from pFP35 decreased sharply at a maximum signal decrease rate 3.5-fold higher than in EXT3 (**Figure 2B, right**). So, only for pFP34 was the composition of NEBExpress favorable for both stages of transcription dynamics, enhancing both Pepper signal increase and stability. The proprietary nature of NEBExpress makes it difficult to reconcile these results. The NEBExpress reaction formulation includes a murine RNase inhibitor, which, at the recommended concentration, improved RNA signal only slightly (**Figure S4A**) and thus could not alone account for this enhancement in transcription dynamics. The same RNase inhibitor had an adverse effect on transcription in EXT3 (**Figure S4B**), although possibly as a result of unsuitable oxidizing conditions in this extract.

We next used templates lacking the *sfGFP* gene or an RBS to measure transcription dynamics in NEBExpress, with results markedly different from those in EXT3. In NEBExpress, removing *sfGFP* improved the maximum Pepper signal and rate of signal increase from both pFP34 and pFP35 (**Figure 2B, center and right)**. However, this modification also resulted in a faster maximum rate of Pepper signal decrease from pFP34, reflecting a higher susceptibility to ribonuclease-mediated mRNA degradation. Removing the RBS from pFP34 also had an adverse effect on template stability, but maintained pFP34’s maximum Pepper signal. For pFP35, removing the RBS was beneficial to all transcription metrics.

We also measured transcription and translation dynamics in PURExpress, a commercially available *E. coli* reconstituted system. Of all CFE systems characterized here, PURExpress stands out, because its mixture of purified proteins does not include ribonucleases and presumably contains a lower concentration of nucleic acids, small molecules, and proteins endogenous to the host organism than extracts. Translation measurements showed that, while PURExpress generated even less sfGFP signal than NEBExpress, PURExpress had the most rapid sfGFP signal increase rate of the three systems (**Figure 2C, left**), a characteristic that makes PURExpress suitable for applications that require rapid CFE performance. Although we added 5 nmol/L of DNA to all CFE systems characterized here to enable a direct comparison, 1 nmol/L plasmid generated more sfGFP than 5 nmol/L in PURExpress (**Figure S5**). With regard to transcription, both the maximum Pepper signal and rate of signal increase were substantially higher in PURExpress than in EXT3 from pFP34 and pFP35 (**Figure 2C, center and right**). The maximum Pepper signals were similar to those in NEBExpress. Interestingly, the Pepper signal in PURExpress did not decay after reaching a maximum value, reflecting the low levels of ribonucleases and indicating that the signal decrease observed in lysates is not inherent to the aptamer. In this ribonuclease-deficient environment, the Pepper signal was still generally higher for pFP34 than pFP35, suggesting that—although to a small extent—tDF30ppr is inherently brighter, folds more efficiently, or binds more effectively to HBC620 than DF30ppr.

Measurements of transcription dynamics in PURExpress with templates lacking a *sfGFP* gene or an RBS contrasted measurements in EXT3 and NEBExpress. In PURExpress, these template modifications had a positive but weak effect on transcription (**Figure 2C, center and right**). These results support our hypothesis that the presence of the RBS and the *sfGFP* gene helped to protect the transcript from degradation by ribonucleases in EXT3. In the ribonuclease-deficient PURExpress, this presumed protection is irrelevant, and the now-improved signal likely results from a diversion of resources from translation to transcription. However, measurements of transcription with templates lacking *sfGFP* or an RBS were not weak in NEBExpress, suggesting other processes may be at play or that NEBExpress confers additional protection against ribonuclease-mediated degradation. Taking together the results for modified templates in three CFE systems, our measurement framework emphasizes the importance of careful nucleic acid template design, in particular for CFE systems susceptible to nuclease-mediated template degradation.

To decouple measurements of translation from transcription, we assessed sfGFP and Pepper expression from mRNA templates generated via *in vitro* transcription from pJL1, pFP34, and pFP35 (**Figure 3**). **Figure 2A** includes translation measurements with 300 nmol/L of mRNA in EXT3, which generated a maximum sfGFP signal similar to the signal from DNA templates, although we also tested other mRNA concentrations (**Figure S6**). Using mRNA instead of DNA templates slightly accelerated the rate of sfGFP signal increase and reduced the time to reach a maximum sfGFP signal in all three CFE systems (**Figure 3A-C, left**), likely because mRNA templates bypass transcription and allow translation to start immediately. Consistent with expression from DNA, transcriptionally fusing *sfGFP* with tDF30ppr reduced sfGFP levels. However, this time, fusing with DF30ppr also had an adverse effect (**Figure 3A-C, left vs. Figure 2A-C, left**).

**Figure 3.**
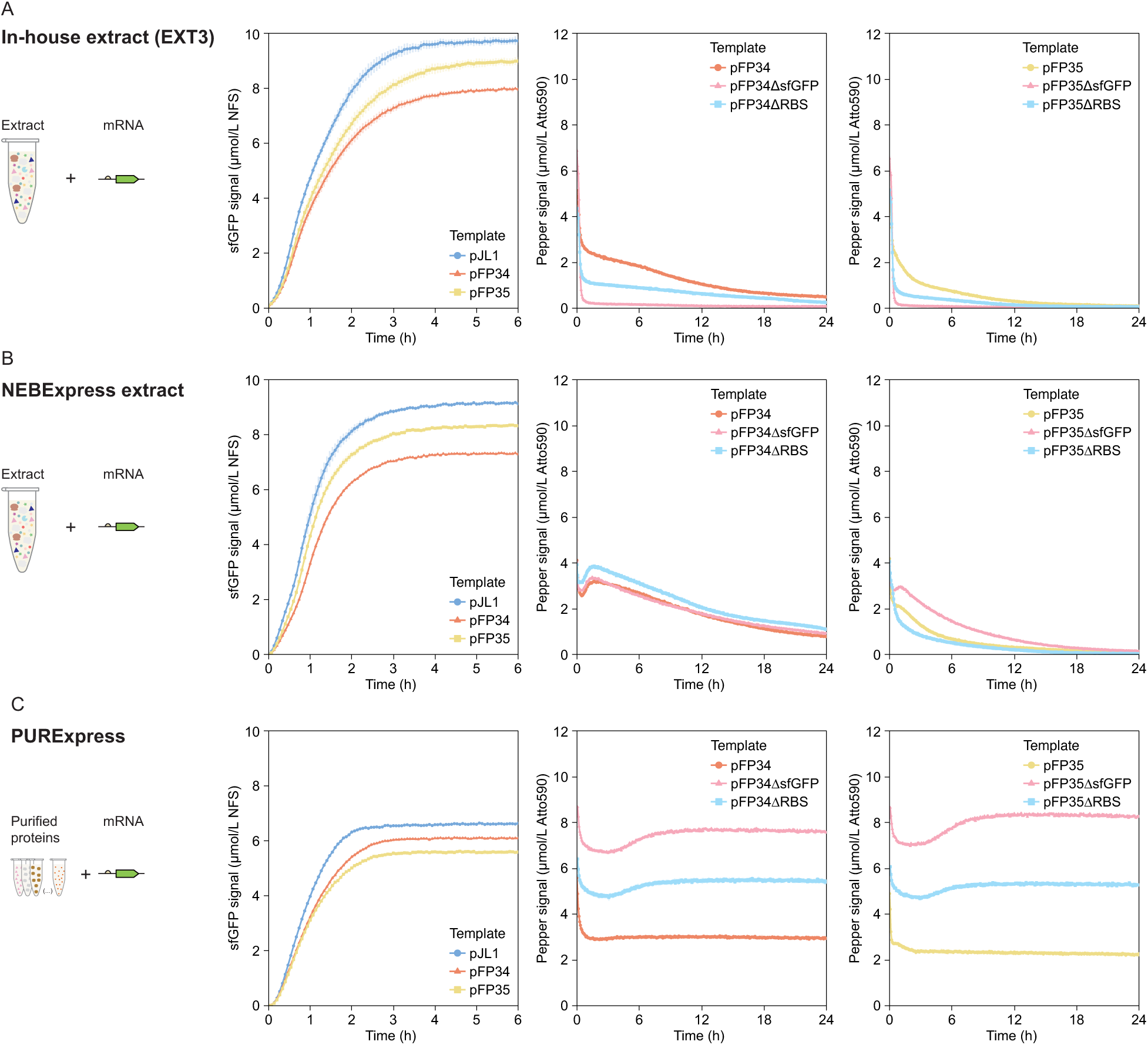
Baseline measurements of transcription and translation dynamics from mRNA templates in **(A)** EXT3, **(B)** NEBExpress, and **(C)** PURExpress. All reactions include mRNA templates added at a concentration of 300 nmol/L. sfGFP measurements are reported in Molecules of Equivalent Soluble Fluorochrome (MESF) of a NIST-traceable fluorescein standard (NFS) and are shown for only the first 6 h of the reaction, because the signal remains constant until 24 h. Pepper mRNA measurements are reported in MESF of Atto 590. The error bars indicate the standard deviation of three technical replicates.

In EXT3, the Pepper signal from mRNA templates during the CFE reaction was highest for the unmodified template, followed by the ΔRBS variant, and then by the ΔsfGFP variant (**Figures 3A, center and right**). For all constructs, the initial Pepper signal decreased sharply in the first 20 min, then decreased steadily at a lower rate. These two apparent signal decrease regimes likely did not represent RNA degradation in CFE systems, as a similar sharp decrease in signal also occurred for mRNAs added to PURExpress, a system with minimal ribonuclease activity (**Figure 3C, center and right**). This decrease in signal at the beginning of the reaction is likely associated with the change in temperature the reaction plate experiences in the pre-warmed plate reader, and is consistent with other experiments, with both DNA and mRNA templates (**Supplementary File 1, Section III**). While this temperature-related effect is common, we did not expect such a substantial decrease in signal and thus hesitate to correlate the initial signal for each template with the strength of transcription from each template. Nonetheless, the similar initial Pepper signal for mRNAs from pFP34 and pFP35 and more rapid signal decrease for mRNA from pFP35 point to a larger difference in stability than in brightness between the two aptamer scaffolds. The continuous decrease in Pepper signal during the CFE reaction is consistent with substantial mRNA degradation. In fact, the rate of Pepper signal decrease from pFP34 mRNA exceeded the maximum rate of Pepper signal decrease from pFP34 DNA in the first 5 h of the CFE reaction with DNA templates, suggesting that processes leading to Pepper signal increase (transcription, folding, and binding to the HBC620 dye) proceeded for several hours. Considering the high mRNA degradation rates, it was surprising that mRNA templates could generate similar sfGFP yields to DNA templates (**Figure 2A vs Figure 3A**); the Pepper signal may not be representative of the number or stability of sfGFP transcripts.

Measurements of transcription dynamics from mRNA templates in NEBExpress captured phenomena we could not capture in EXT3 and helped to elucidate differences between the two extracts. In NEBExpress, the initial decrease in Pepper signal was not as substantial as in EXT3 (**Figure 3B-C, center and right**). A short-lived signal increase followed this initial signal decrease, likely reflecting binding of the Pepper mRNA to the HBC620 dye; in EXT3, ribonuclease-mediated mRNA degradation likely exceeded the fluorescence signal produced by this binding. The Pepper signal maintained higher values in NEBExpress than in EXT3, although this difference in signal cannot account for the pronounced difference observed with DNA templates (**Figure 3B vs. Figure 2B**). Instead, this result suggests that the high Pepper signals observed with DNA templates in NEBExpress resulted mostly from stronger transcription rather than enhanced aptamer folding, binding to the dye, or attributes of the dye. The Pepper signal from mRNAs templates lacking *sfGFP* further supports this idea: the signal from the pFP34ΔsfGFP mRNA was similar to the signal from pFP34 mRNA, despite markedly higher signal from pFP34ΔsfGFP DNA than its unmodified counterpart.

Measurements with mRNA templates in PUREexpress exhibited different trends from mRNAs compared with EXT3. While the Pepper signal from mRNAs in PURExpress also underwent an initial decrease, the signal slightly recovered and remained stable, reflecting the low levels of ribonucleases (**Figure 3C, center and right**). At higher mRNA concentrations, the Pepper signal increased immediately, likely as a result of Pepper binding to the HBC620 dye, before decreasing to a stable level (data not shown); this behavior is similar to that observed in NEBExpress. Consistent with expression from DNA templates, the Pepper signal from mRNA templates was highest for the ΔsfGFP variant, followed by the ΔRBS variant, and then by the unmodified template—opposite to the trend in EXT3. In addition, both sfGFP and Pepper signals from tDF30ppr-encoding mRNAs were again only slightly higher than the signals from DF30ppr-encoding counterparts, a difference more pronounced at lower mRNA concentrations (**Figure S7**).

### Probing transcription and translation dynamics of different CFE systems

To assess the ability of our transcription and translation measurements to resolve differences between CFE systems, we prepared a suite of extracts with deliberate modifications to either the host strain or the preparation conditions (**Figure 4A**). These modifications are primarily known to affect protein production in T7 RNAP-based CFE systems, but their effects on transcription are not as clear in literature. For these extracts, we do not show results for pJL1, because pJL1 exhibited similar translation dynamics to pFP34 and pFP35 in EXT3.

**Figure 4.**
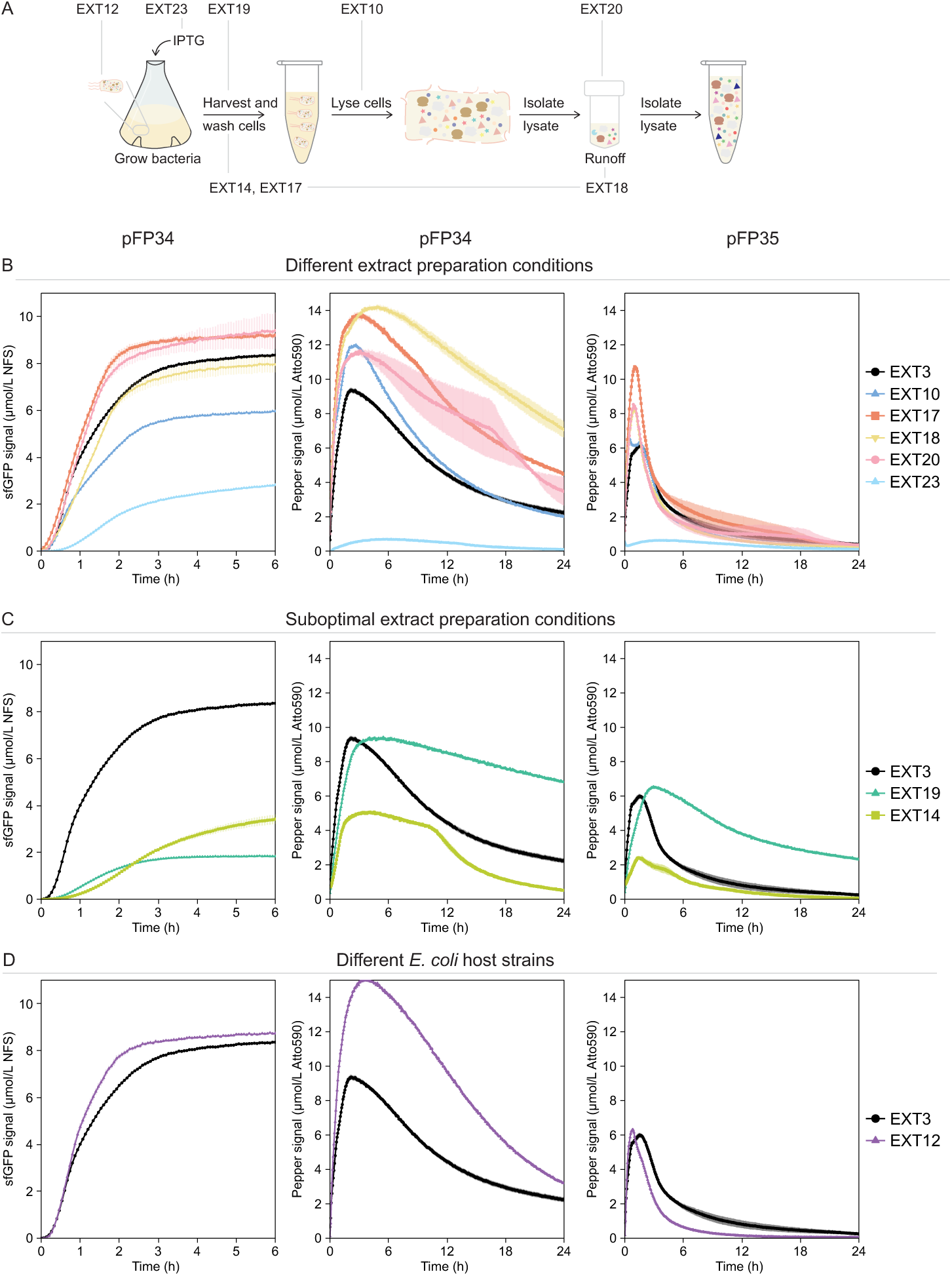
Measurements of transcription and translation dynamics in extracts prepared in house **(B)** under different conditions, **(C)** under suboptimal conditions, and **(D)** from different *E. coli* host strains. Panel A specifies the step of extract preparation modified for each extract. Panels B-D include data for sfGFP signal from pFP34 (left), Pepper signal from pFP34 (center), and Pepper signal from pFP35 (right), with the relevant DNA template added at 5 nmol/L. sfGFP expression from pFP35, due to its similarity to expression from pFP34, is shown in **Figures S8**. sfGFP measurements are reported in Molecules of Equivalent Soluble Fluorochrome (MESF) for a NIST-traceable fluorescein standard (NFS) and are shown for only the first 6 h of the reaction because the signal remains constant until 24 h. Pepper mRNA measurements are reported in MESF for Atto 590. The error bars indicate the standard deviation of three technical replicates.

We first characterized CFE systems from BL21 Star (DE3) extracts prepared under different conditions (**Figure 4B**). EXT10 had the same preparation conditions as EXT3 except for the lysis step: instead of a single-tip sonicator, it was lysed with a 24-tip sonicator, which, although adjusted to the same settings, lacked control over the energy delivered by each tip. The sonicator reports a total energy input equal to the energy generated at the transducer, not each tip; based on this number and assuming each tip exerts an equal amount of energy, each tip exerted a slightly lower sonication energy than when using a single tip. Although the energy input applied to disrupt cells affects CFE in ways we do not yet fully understand(*43*), lower sonication energy input typically reduces the extract’s endogenous protein content and adversely affects T7 RNAP-driven CFE(*25, 43*). Inconsistent with this expectation, EXT10 and EXT3 had statistically identical protein contents based on a Bradford assay, so any change in performance could not be attributed to a change in total protein levels (**Figure S2**). However, with regard to translation, the maximum sfGFP signal and signal increase rate in EXT10 were, respectively, about 40 % and 50 % lower than in EXT3. Surprisingly, this change in sonication conditions had a positive effect on transcription, improving the maximum Pepper signal and signal increase rate from both pFP34 (**Figure 4B, center**) and pFP35 (**Figure 4B, right**).

Unlike EXT3, EXT20 did not undergo a runoff reaction, which is a post-lysis incubation at 37 °C to promote degradation of genomic material, release ribosomes from mRNAs, and improve transcription from endogenous promoters(*32*). In EXT20, the maximum rate of sfGFP signal increase was slightly higher than in EXT3, and the signal sustained higher rates until reaching a maximum value 9.0 % higher. Omitting the runoff reaction had more substantial effects on transcription, improving the maximum Pepper signal and signal increase rate by 24 % and 85 %, respectively, for pFP34 (**Figure 4B, center**), and 39 % and 59 %, respectively, for pFP35 (**Figure 4B, right**). Without a runoff reaction, EXT20 was not exposed to 37 °C and likely retained a higher concentration of endogenous nucleic acids, which can have a positive effect on target gene expression(*43, 44*). In addition, the runoff reaction adversely affects T7 RNAP-driven sfGFP expression, and the strength of this effect varies with the host organism or even strain(*25, 32*).

EXT17 used a glutamate-based buffer instead of EXT3’s acetate-based buffer to wash cells after harvest, a substitution reported to improve protein yield(*30, 31*). The change in buffer had a minimal effect on translation, resulting in 11 % higher maximum sfGFP signal, but a 6.0 % lower maximum sfGFP signal increase rate (**Figure 4A, left**). Again, the effect on transcription was more substantial: the glutamate-based buffer improved the maximum Pepper signal and increase rate by 47 % and 65 %, respectively, for pFP34 (**Figure 4A, center**), and 75 % and 32 %, respectively, for pFP35 (**Figure 4A, right**).

EXT18 also used a glutamate-based buffer and did not undergo a runoff reaction. Individually (EXT20 and EXT17), these changes to preparation conditions were beneficial to transcription and had a minor effect on translation relative to EXT3. Together, they still minimally affected translation, but, with pFP34, they improved the maximum Pepper signal and signal increase rate by 52 % and 42 %, respectively, relative to EXT3 (**Figure 4A, center**). With pFP35, however, we measured a 33 % higher maximum Pepper signal but a 14 % lower maximum signal increase rate in EXT18 (**Figure 4A, right**). Of all extracts, EXT18 exhibited the highest Pepper signal value after 24 h, which resulted not from a lower rate of signal decrease but from a broad peak in signal.

EXT23 was prepared from cells not supplemented with IPTG to induce expression of T7 RNAP, a protocol modification useful in applications that use endogenous promoters or do not require maximizing protein levels(*45*). With regard to translation, omitting IPTG reduced the maximum sfGFP signal increase rate 5-fold and the maximum sfGFP signal 2.7-fold compared with EXT3 (**Figure 4A, left**). This modification had a more substantial adverse effect on transcription: with pFP34, for example, it caused a 14-fold reduction in the maximum Pepper signal and a 34-fold reduction in the maximum signal increase rate (**Figure 4A, center**). These results show a tradeoff between transcription signal strength and longevity. While the Pepper signal for pFP34 and pFP35 continued to increase for another 3.7 h and 2 h, respectively, both the maximum Pepper signal and signal increase rate were lower in EXT23 than in EXT3.

Applying our measurement framework to different extracts showed that quantitative characterization of CFE systems depends on the specific measurement tool used to conduct the measurement. Modifications to extract preparation conditions affected expression from pFP34 and pFP35 differently. Translation measurements exhibited highest maximum sfGFP signals, highest maximum rates of signal increase, and lowest maximum rates of signal decrease in EXT17 and EXT20 with both pFP34 and pFP35. However, transcription measurements with pFP34 exhibited optimal reaction metrics in a different extract than with pFP35. Whereas the maximum Pepper signal from pFP34 was highest in EXT18, the signal from pFP35 was only the third highest maximum Pepper signal in that extract. In addition, certain modifications to extract preparation conditions improved all reaction metrics but to different extents depending on the DNA template. For example, modifying EXT3 to generate EXT17 enhanced the maximum Pepper signal from pFP35 to a greater extent than the maximum rate of signal increase; for pFP34, the maximum rate improved more than the maximum signal. We also characterized all in-house extracts using modified DNA templates lacking the *sfGFP* gene or the RBS, and found that the Pepper signal in these extracts was generally higher with modified templates than with unmodified counterparts, opposing the results observed in EXT3 (**Figure S9**). Strategies to tune a CFE system towards a specific application may enhance one metric but worsen another, and one extract may appear more or less productive than another depending on the nucleic acid template used as a measurement tool.

Having primarily explored changes to extract preparation that improved performance and seeking to elucidate differences between high- and low-performing systems, we deliberately generated low-performing extracts by selecting suboptimal preparation conditions, starting with changes to the cell harvest time. Historically, lysate preparation protocols recommend harvesting cells in mid-exponential growth phase—equivalent to (1.5 to 1.8) OD600 for *E. coli* BL21 Star (DE3) growth in 2xYTP—, when cellular metabolism is most active and protein expression machinery is present at high concentrations. However, Failmezger *et al*. found that *E. coli* extracts from cells harvested in stationary phase had a similar eGFP yield to an extract from mid-exponential phase cells(*46*), challenging well-established protocols. Measurements of translation dynamics in extracts from cells at OD600 2.0 and 2.2 (EXT15 and EXT16, respectively) exhibited lower maximum sfGFP signals and signal increase rates, but higher maximum Pepper signals than EXT3 (**Figure S10**). We prepared an extract (EXT19) with a substantial—4.5-fold—reduction in sfGFP signal only after allowing cells to grow for 24 h prior to harvest (**Figure 4B, left**). Translation measurements in EXT19 reached this substantially lower maximum sfGFP signal at a 6.7-fold lower maximum signal increase rate than in EXT3, but 20 min earlier than in EXT3, indicating that weaker protein production allowed enhanced reaction longevity. Poor translation measurements in EXT19 did not extend to transcription. In fact, the maximum Pepper signals from pFP34 were statistically identical in EXT3 and EXT19 (**Figure 4B, center**), although this signal occurred 3.0 h later and at a lower rate in EXT19. The maximum Pepper signal from pFP35 was higher in EXT19 than in EXT3, but, similar to pFP34, occurred over 2 h later and at a lower rate (**Figure 4B, right**). Notably, the Pepper signals at 24 h from both pFP34 and pFP35 were higher—3-fold and 8.9-fold, respectively—than in EXT3, reflecting much lower signal decrease rates in EXT19. Perhaps EXT19 could sustain transcription for longer or degrade RNAs at a lower rate. EXT19 may be deficient in key ribonucleases that are depleted or less efficient during stationary cellular growth(*47, 48*). Whether a product of improved transcription or diminished degradation, this increase in the amount and duration of Pepper signal did not correspond to enhanced sfGFP production, although EXT19 could have become saturated with Pepper transcripts, preventing further protein production. The strong Pepper signals and weak sfGFP signals in EXT19 are consistent with previous reports of a tradeoff between transcription and translation(*44, 49*).

To characterize another low-performing system, we resuspended the cells used to prepare EXT14 with a buffer at a pH of 10, higher than the recommended 7.7 and likely to damage endogenous proteins and nucleic acids. Both transcription and translation dynamics were weaker in EXT14 than in EXT3. The maximum sfGFP signal from pFP34 was 2.3-fold lower and reached over 2 h later in EXT14, indicating an apparent tradeoff between protein yield and reaction longevity unlike what we observed in EXT19 (**Figure 4B, left**). Consistent with the other low-performing extract (EXT19), measurements of transcription dynamics were enhanced in EXT14 relative to EXT3, but exhibited lower rates of signal decrease (**Figure 4B, center and right**). Interestingly, the peak Pepper signal from pFP34 remained nearly constant for several h before decreasing, pointing to similar Pepper signal increase rates and RNA degradation rates for several h during the CFE reaction (**Figure 4B, center**).

To study the effect of a different *E. coli* host strain, we characterized EXT12, an extract identical to EXT3 in preparation conditions but derived from an *E. coli* BL21 (DE3) strain with an intact RNase E and thus presumably lower RNA stability. The identity of the host strain had a slight effect on translation, with higher maximum sfGFP signal and maximum rate of signal increase in EXT12 than in EXT3 (**Figure 4C, left**). The effect of an intact RNase E on transcription was more pronounced, in particular, with pFP34 as the template. In EXT12, measurements of transcription with pFP34 had an 80 % faster maximum rate of Pepper signal increase and reached a 61 % higher maximum Pepper signal 92 min later than in EXT3, thus exhibiting faster and longer duration of signal increase (**Figure 4C, center**). While we expected RNA degradation to be stronger in EXT12, we measured a maximum rate of Pepper signal decrease only 4 % lower than in EXT3, within experimental standard deviation. With pFP35, the maximum Pepper signal and rate of signal increase were also higher in EXT12 than in EXT3 (**Figure 4C, right**). However, consistent with the expectation of enhanced RNA degradation in EXT12, the maximum rate of Pepper signal decrease exceeded the rate in EXT3 by 31 %. It is unclear why the change in transcription dynamics in EXT12 varied so much across Pepper aptamer scaffolds; these results support the previous observation that the measured system performance depends on the measurement tool.

To expand the utility and assess the sensitivity of our measurement framework beyond T7 RNAP-driven transcription, we included measurements for *E. coli* RNAP-driven transcription. Because T7 RNAP catalyzes transcription more efficiently than *E. coli* RNAP and does not target promoter sequences endogenous to *E. coli*, T7 RNAP has become the workhouse for high-yield recombinant protein expression in *E. coli*(*24, 50, 51*). T7 RNAP is a single-unit polymerase less susceptible to global gene regulation in BL21 (DE3), whereas *E. coli* RNAP has 12 units that must be expressed and assembled to generate a functional polymerase. Considering these features and that T7 RNAP is typically overexpressed in BL21 (DE3) prior to extract preparation, T7 RNAP-driven expression in extracts is robust and less sensitive to differences among CFE systems. Therefore, *E. coli* RNAP-driven expression could provide additional insight. Because we expect *E. coli* RNAP-driven transcription to be weak, probing this transcription also assesses the sensitivity of our framework’s measurements of transcription dynamics. Moreover, an increasing number of CFE applications, such as biosensing(*45, 52, 53*), involve genetic circuits with endogenous promoters. To probe CFE driven by *E. coli* RNAP, we replaced the T7 promoter with a strong *E. coli* σ^70^ promoter, P_J23100_, in pJL1, pFP34, and pFP35 to generate, respectively, pFP24, pFP59, and pFP60 (**Figure 5A**). We then used these templates to measure transcription and translation dynamics in our extracts prepared in house (**Table 1**).

**Figure 5.**
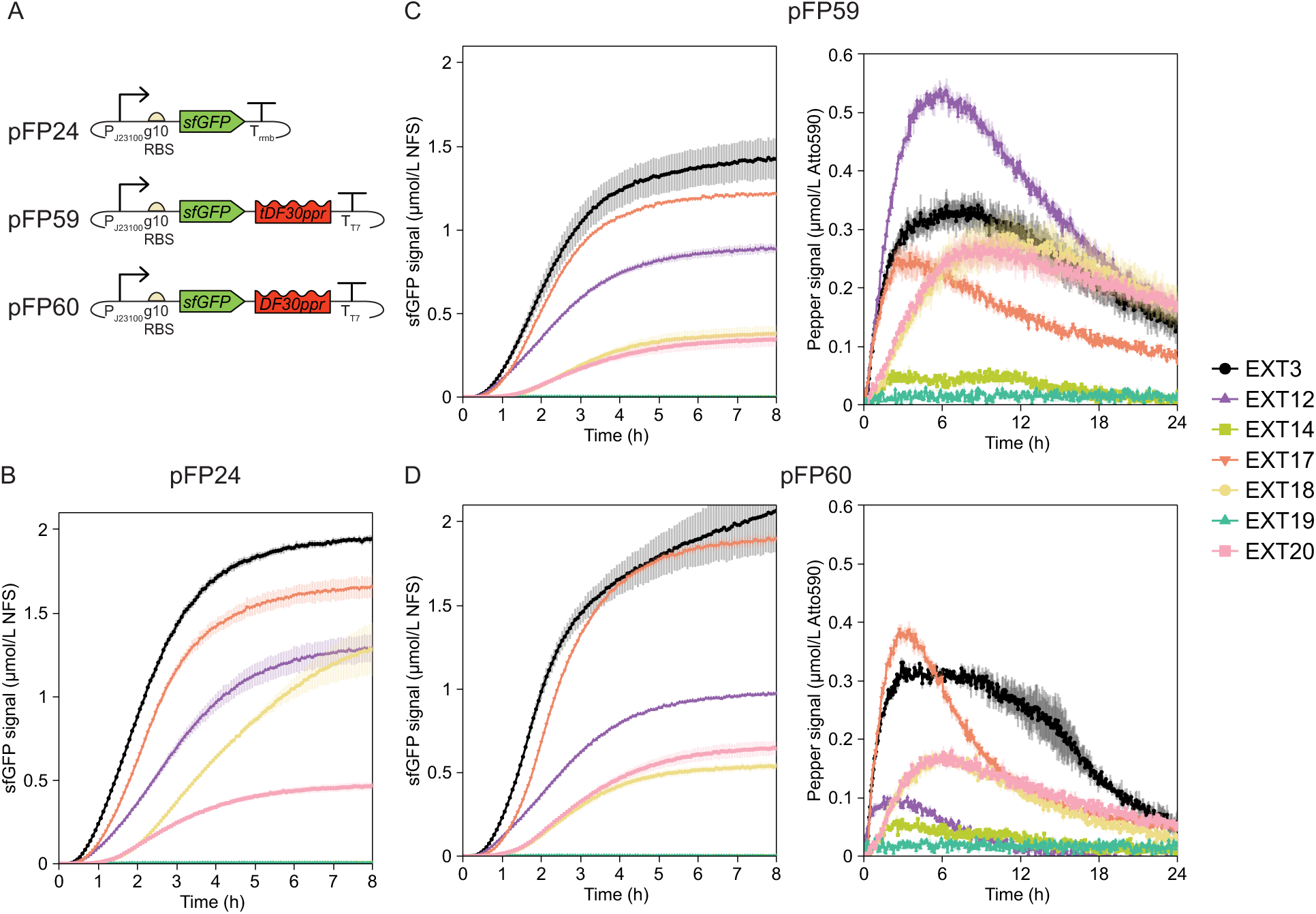
Measurements of transcription and translation dynamics using DNA templates encoding *E. coli* RNAP-driven transcription. **(A)** Plasmids pFP24, pFP59, and pFP60 are, respectively, based on pJL1, pFP34, and pFP35, but, instead of T7 phage transcription elements, they contain a promoter (P_J23100_) recognized by *E. coli*’s RNAP and housekeeping sigma factor (σ^70^). **(B)** Measurements of translation from pFP24. **(C)** Measurements of translation (left) and transcription (right) dynamics from pFP59. **(D)** Measurements of translation (left) and transcription (right) dynamics from pFP60. sfGFP measurements are reported in Molecules of Equivalent Soluble Fluorochrome (MESF) for a NIST-traceable fluorescein standard (NFS) and are shown for only the first 8 h of the reaction to highlight differences among extracts. Pepper mRNA measurements are reported in MESF for Atto 590. The scale of the y-axis was adjusted due to lower expression levels from endogenous transcription machinery relative to T7 RNAP-driven transcription. The error bars indicate the standard deviation of three technical replicates.

Transcription and translation were less efficient for the templates with P_J23100_ instead of P_T7_. The differences in transcription were more substantial: in EXT3, the maximum Pepper signal was 25-fold lower for pFP59 (**Figure 4B, center vs. Figure 5C, right**), and 19-fold fold lower for pFP60 (**Figure 4B, right vs. Figure 5D, right**) compared with their P_T7_ counterparts. Similarly, translation measurements showed lower sfGFP signals, with P_J23100_ templates generating around 4-fold less sfGFP in EXT3 (**Figures 4A-C**). In all extracts but EXT3, both transcription and translation from P_J23100_ templates had a lag during which the signal was indistinguishable from background levels, reflecting slower *E. coli* RNAP-catalyzed transcription. Our measurement framework could measure transcription and translation dynamics with the relatively weak P_J23100_, but may not be suitable for characterization with even weaker promoters; higher DNA concentrations could alleviate detection issues (**Figure S11**).

Although P_J23100_ templates exhibited optimal translation metrics in EXT3 out of all extracts, the relative performance of the other extracts varied with the scaffold of the Pepper aptamer. Except for expression from pFP24 in EXT18, the sfGFP signal followed the same trend across extracts for all three P_J23100_ templates. In EXT3, measurements of transcription with pFP59 had a higher rate of Pepper signal increase and a lower rate of Pepper signal decrease than with pFP60, but the maximum Pepper signals were identical for pFP59 and pFP60 (**Figure 4C, right vs. Figure 4D, right**), at odds with the marked differences between Pepper aptamers in other extracts. In addition, although the sfGFP signals from pFP59 and pFP60 were similar in EXT12, transcription measurements were strikingly different: the maximum Pepper signal and rate of signal increase from pFP59 were maximal in EXT12 (**Figure 5C, right**), whereas transcription from pFP60 was only better in EXT12 than in the low-performing extracts, EXT14 and EXT19 (**Figure 5D, right**).

Measurements of *E. coli* RNAP-driven CFE differed from T7 RNAP-based measurements but could complement those measurements and help ascertain differences among systems with greater resolution. For example, P_J23100_ templates had inefficient transcription and translation in EXT20, whereas P_T7_ templates yielded better reaction metrics in EXT20 than in EXT3 (**Figure 4A**), consistent with reports of runoff reactions enhancing *E. coli* RNAP-mediated but not T7 RNAP-mediated CFE(*32*). In extracts prepared under suboptimal conditions, measurements with P_T7_ templates exhibited better (EXT19) or slightly worse (EXT14) transcription metrics than in EXT3 despite poor yet measurable sfGFP signal (**Figure 4B**). However, measurements with P_J23100_ templates had low Pepper and sfGFP signals in EXT19 and EXT14 (**Figures 5C, 5D**). A glutamate-based buffer (EXT17 and EXT18) was beneficial to T7 RNAP-mediated transcription but not to *E. coli* RNAP-mediated transcription (**Figure 4A vs. Figures 5C-D**). Taken together, these results suggest that characterization of CFE systems via T7 RNAP-based measurements may not be relevant to applications that harness *E. coli* RNAP for CFE and *vice-versa*. Characterization efforts, therefore, should include application-specific measurement tools and metrics.

After characterizing different extracts and transcription machineries, we measured the effect of reaction volume and vessel, exploring reaction formats beyond 10 µL volumes in a clear 384-well plate. Specifically, we included measurements for two additional reaction volumes—5 and 20 µL—and a black, flat-bottom 384-well plate with larger wells. Black plates are also commonly used for CFE, in particular for fluorescence measurements due to lower autofluorescence than clear plates. To compare results across microplates and volumes, we generated calibration curves for the two fluorochromes in each vessel and at each volume (**Figure S1**).

With regard to translation, the maximum sfGFP signal and rate of signal increase were inversely proportional to the reaction volume for both pFP34 and pFP35 in both plates (**Figures 6A-B, left**). This trend is likely related to the reduced availability of oxygen in the headspace of the well with increasing reaction volume, which is detrimental to transcription and translation due to oxygen’s role in ATP production via oxidative phosphorylation(*15*). The adverse effect of increasing the reaction volume was less pronounced in the black plate with larger wells and thus larger headspace. Also potentially related to its larger headspace, the black plate had a higher maximum sfGFP signal at a given volume than the clear plate, a difference more pronounced at lower volumes. At lower volumes in both plates, despite the faster maximum sfGFP signal increase rate, reactions took longer to reach the maximum sfGFP signal, inconsistent with previous reports of a tradeoff between reaction rate and longevity(*54, 55*).

**Figure 6.**
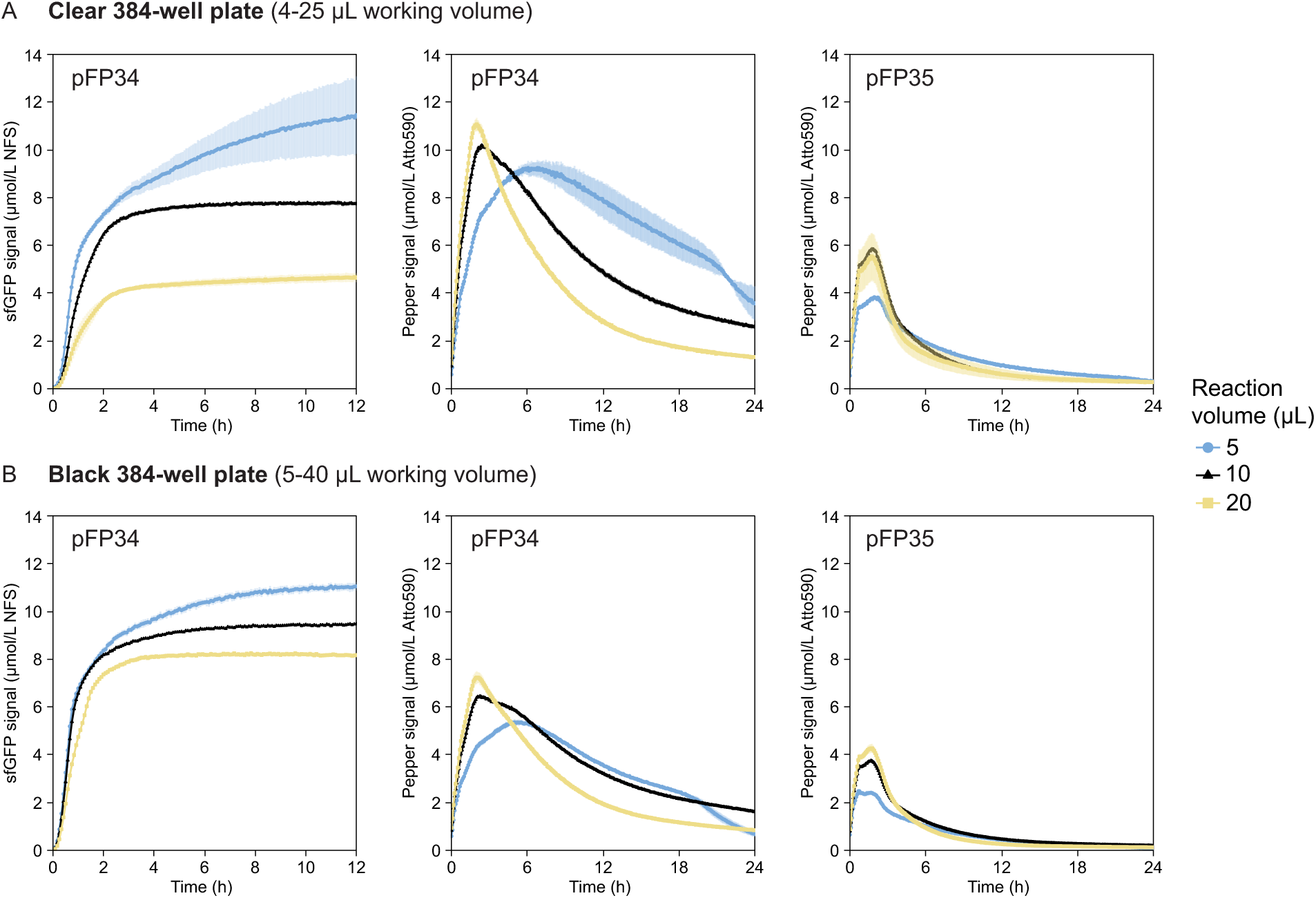
Measurements of transcription and translation dynamics in different reaction vessels and volumes in EXT3. **(A)** (5, 10, 20) µL reactions in a clear, flat-bottom 384-well plate with a working volume of (4 to 25) µL. **(B)** (5, 10, 20) µL reactions in a black, flat-bottom 384-well plate with a working volume of (5 to 40) µL. sfGFP expression from pFP35, due to its similarity to expression from pFP34, is shown in **Figures S8B.** sfGFP measurements are reported in Molecules of Equivalent Soluble Fluorochrome (MESF) of a NIST-traceable fluorescein standard (NFS) and are shown for the first 12 h of the reaction, instead of the 6 h used in previous Figures, because low-volume reactions had longer lifetimes. Pepper mRNA measurements are reported in MESF of Atto 590. The error bars indicate the standard deviation of three technical replicates.

Contrary to the trends in translation, measurements of transcription showed higher maximum Pepper signals and rates of signal increase at larger reaction volumes (**Figures 6A-B, center and right**). However, larger-volume reactions also had higher maximum rates of Pepper signal decrease, indicating either more efficient RNA degradation or less efficient processes leading to signal increase—aptamer transcription, folding, and binding to HBC620. At a given volume, the clear plate had higher Pepper signals than the black plate, perhaps due to stronger autofluorescence. These trends did not vary with the Pepper aptamer scaffold.

Measurements in different reaction formats showed that calibration curves matching reaction specifications are essential to preventing misinterpretation of assay results. In the black plate, sfGFP fluorescence intensity values (in arbitrary units) were (3 to 7)-fold higher than in the clear plate and had a directly proportional relationship with the reaction volume (**Figure S12**), suggesting that reactions in the black plate generated more sfGFP and benefitted from larger volumes. However, upon calibrating fluorescence values, the sfGFP signal was at most 1.9-fold higher in the black plate than in the clear plate, and was higher at lower volumes (**Figure 6B, left**). These differences may be exacerbated at larger scales and call for careful analysis of CFE data.

Lastly, we assessed the ability of our measurements to characterize CFE systems supplemented with energy systems that use different molecules—PEP or pyruvate—to regenerate ATP and fuel transcription and translation. The PANOx-SP energy system used in this work, which contains PEP but not pyruvate, provides all the additional molecules (e.g., NAD and CoA) required to regenerate ATP from PEP and pyruvate and to harness pyruvate as a secondary energy source to PEP(*56*). Of all the formulations tested, the PANOx-SP system yielded the best transcription and translation metrics (**Figure 7**). In the absence of both PEP and pyruvate, both processes were inefficient but could still proceed, likely because other small molecules included in the reagent mix, such as magnesium glutamate, could help sustain these processes. Using only pyruvate had a more pronounced effect on translation than on transcription, producing a Pepper signal profile resembling a reaction lacking both PEP and pyruvate (**Figure 7A-B, right**). Combining PEP with pyruvate reduced the maximum sfGFP signal slightly for pFP34 and pFP35 (**Figure 7A-B, left**). Achieving improved transcription and translation signals in CFE systems supplemented with PEP instead of pyruvate is not surprising, as PEP conversion to pyruvate regenerates an extra molecule of ATP compared to pyruvate. However, the diminishing returns observed in CFE systems including both PEP and pyruvate contradict a previous report of a synergistic effect between the two molecules(*56*). Our measurement framework captured transcription and translation nuances in CFE systems with different energy sources and could be valuable for evaluating other energy systems.

**Figure 7.**
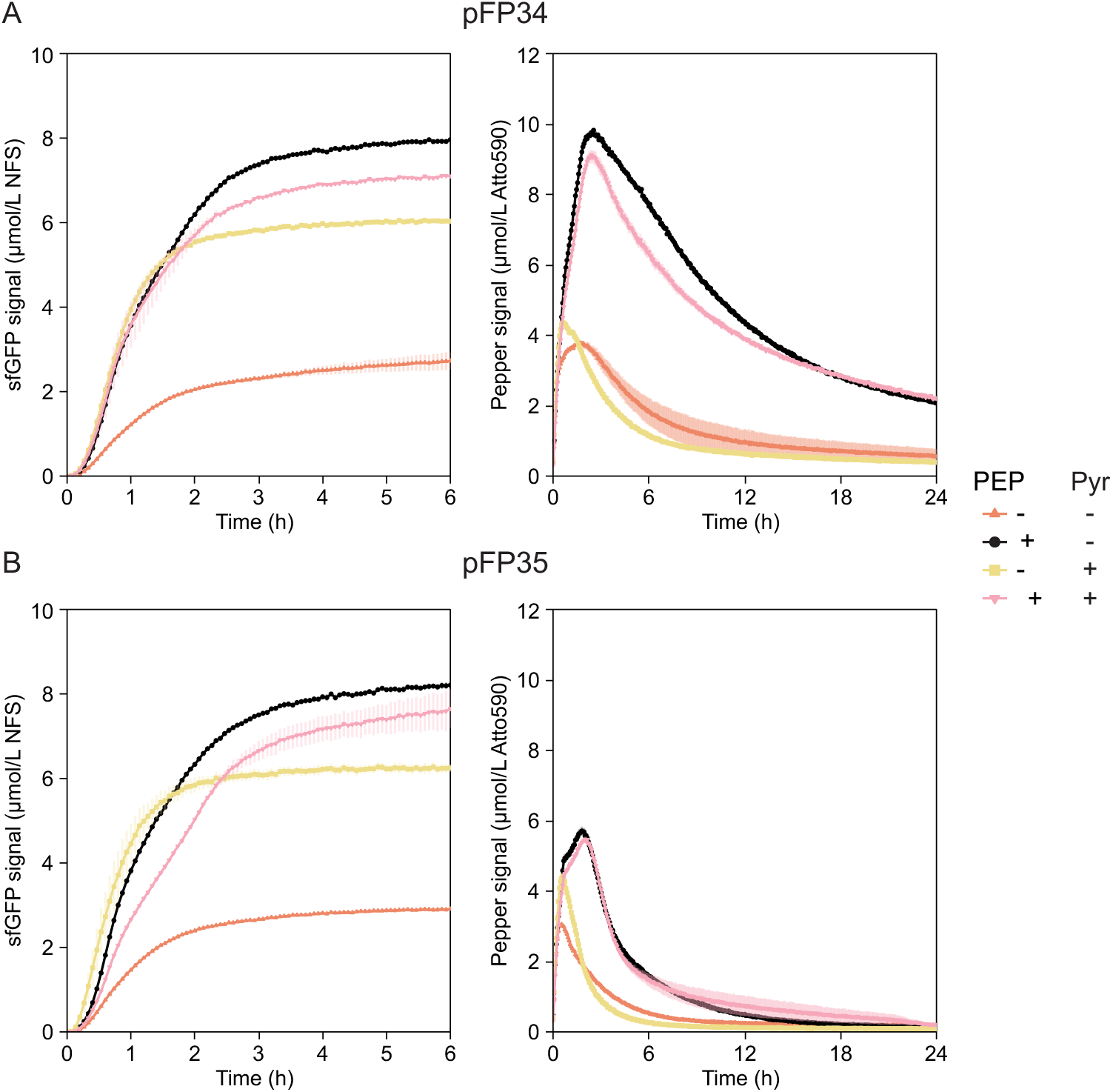
Measurements of transcription and translation dynamics in EXT3 supplemented with different energy substrates phosphoenolpyruvate (PEP) or pyruvate for ATP regeneration for expression from **(A)** pFP34 and **(B)** pFP35. In each panel, the left plot shows translation measurements and the right plot shows transcription measurements. sfGFP measurements are reported in Molecules of Equivalent Soluble Fluorochrome (MESF) for a NIST-traceable fluorescein standard (NFS) and are shown for only the first 6 h of the reaction because the signal remained constant until 24 h. Pepper mRNA measurements are reported in MESF for Atto 590. The error bars indicate the standard deviation of three technical replicates.

## Discussion

Including measurements of transcription dynamics challenged our perception of CFE system performance and provided additional insight into how transcription and translation correlate. Measurements of transcription dynamics were more sensitive to changes to extract preparation, and thus facilitated comparison of systems that did not seem to differ considerably based on their similar sfGFP signals. In addition, certain systems with low sfGFP signals, which would typically be classified as low-performing, exhibited high Pepper signals and could be useful in RNA-based applications. With measurements of both sfGFP and Pepper signals, we observed more nuanced effects of different extract preparations on both transcription and translation. These two processes share a complex relationship, as translation follows transcription and uses similar resources, usually resulting in a tradeoff in efficiency between the two reactions. Certain modifications to extract preparation, such as in EXT10, caused such a tradeoff, leading to weaker sfGFP signals but stronger Pepper signals than in EXT3. However, other modifications improved or weakened both signals, pointing to a mechanism that affects global performance, such as an increase or decrease in the availability of resources shared by transcription and translation. The two measurements also probed transcription and translation longevity, showing that transcription occurred rapidly before RNA degradation outpaced it, and translation proceeded for several additional hours.

Measurements with RNA templates supplemented our understanding of where differences among systems arise. Importantly, these measurements were useful for decoupling aptamer transcription from aptamer folding, binding to dye, and degradation. For example, measurements of Pepper signal with mRNA templates were similar in EXT3 and commercial CFE systems, suggesting that the substantial difference between the Pepper signal generated with DNA templates resulted primarily from a difference in transcription efficiency.

While nucleic acid templates distinguished CFE systems on the basis of transcription and translation under a variety of conditions, they had limitations as measurement tools. Components of a nucleic acid template affected one another, with, for example, removal of the RBS resulting in lower Pepper signal in EXT3 (**Figures 2-3**) but higher Pepper signal in other in-house extracts (**Figure S9**). For extracts prepared in house, system performance depended on which Pepper aptamer was used to measure transcription dynamics, thereby misleading characterization efforts. Furthermore, the quality—based on breakage and purity—of nucleic acid templates affects CFE(*57, 58*) and could interfere with the reproducibility and sensitivity of our assays.

Fluorogenic Pepper RNA aptamers are a valuable tool for measuring transcription dynamics in CFE systems. In particular, they are useful for capturing differences among CFE systems that appeared similar based on the sfGFP signal, and for unveiling the potential value of systems with poor sfGFP signal but strong Pepper signal. Whereas sfGFP is a stable protein, RNA aptamers vary in stability depending on ion levels, additional scaffolds, and other components of the nucleic acid template encoding the aptamer. This sensitivity enables aptamers to capture subtle changes in a system and confers them a modularity that we can leverage to facilitate measurements. For example, we could improve the stability of an RNA aptamer to enable measurements even in low-performing systems.

However, fluorogenic RNA aptamers also pose measurement challenges. The same sensitivity that makes RNA aptamers modular measurement tools also confounds measurements. The dependence of the RNA aptamer signal on other components of the aptamer’s nucleic acid template constrains the design of the nucleic acid template encoding the aptamer, especially in lysate-based CFE systems. In addition, RNA aptamers are susceptible to degradation by ribonucleases in CFE systems, masking the effect of uneven transcription and translation resource allocation, and in certain cases, precluding measurements at later time points of a CFE reaction. The instability of RNAs in extracts also hinders the development of meaningful calibration curves. Different approaches to this issue have been reported (*44, 59–61*), with no consensus achieved to date. Importantly, RNA aptamers do not provide a reliable metric of transcription of the gene with which they are transcriptionally fused. Overall, existing aptamers are not fit for purpose for quantitative measurements in CFE reactions, as they were originally developed for use in cells, often as a qualitative metric. We need a reliable way to monitor transcription dynamics, as alternative tools to RNA aptamers also have limitations(*62–64*).

Our work highlights the importance of calibrating fluorescence intensity data to absolute units. The fluorochromes used in this work enable comparison of fluorescence data collected in different days and different instruments. Notably, the NIST fluorescein standard is directly traceable to a standard reference material. Atto 590, however, is not a standard reference material and does not exactly match the wavelengths of the excitation and emission peaks of the HBC620-Pepper complex. In addition, fluorescein-based calibration is not sufficient for applications involving precise control and measurement of protein or RNA yield or data collection for computational models. Calibration curves based on purified reporters (sfGFP and Pepper) would have been more suitable for these applications and more directly useful to the community. However, we refrained from using purified protein-based calibration because protein purification protocols differ across laboratories, and these reporters are not commercially available in purified form. A calibration curve based on purified Pepper mRNA would have also been difficult to develop due to mRNA degradation in extracts over the course of a CFE reaction. Whether based on dyes or purified proteins, calibration curves should be generated routinely for each reaction format to prevent erroneous data interpretation.

## Conclusions

Here, we demonstrate a measurement framework to characterize transcription and translation dynamics in CFE systems, including a suite of assays using nucleic acid templates and methods for implementation. Our nucleic acid-based assays generate measurements for multiple CFE systems that can serve as an internal reference for users who make their own extracts or purchase commercially available CFE systems. We expand upon measurements of sfGFP fluorescence at a single time point to include time-course data for both sfGFP and Pepper mRNA, providing more detailed characterization of CFE for applications beyond protein synthesis. From these measurements, we compute quantitative metrics that measure the effect of changing the preparation conditions of extracts, and the vessel, volume, and formulation of CFE reactions. We show that the components of the nucleic acid templates can bias measurements and call for the development of fit-for-purpose tools for the development and routine characterization of CFE systems. In testing different reaction formats, we demonstrate the importance of calibration to enable comparison across reaction formats and ensure accurate interpretation of experimental results. With rigorously calibrated data, we can now begin to discuss and share data collected in different laboratories meaningfully.

This work addresses several outstanding measurement challenges that can be partly traced back to insufficient characterization of CFE and hold back applications and adoption of CFE systems. Notably, system performance varies across laboratories and batches(*32, 65, 66*), and even across lots of commercial reconstituted systems(*41*), despite efforts to improve reproducibility(*66*). In addition, published protocols for preparing and using CFE systems, protocol nuances, reagent grades, and reaction assembly techniques remain poorly documented and understood, exacerbating reproducibility issues(*67, 68*). Additional characterization of CFE systems can help identify factors leading to reproducibility issues and protocol details that should be documented routinely. Our limited understanding of CFE systems has also hindered modeling efforts, scale-up, technology transfer(*67*), and the development of fit-for-purpose standards, metrics, and best practices necessary to advance these systems.

Our measurement framework provides a global assessment of system performance based on transcription and translation dynamics that can complement more targeted characterization efforts. As the application repertoire of CFE systems expands, targeted measurements of a subset of molecules relevant to specific applications become increasingly important. Commercially available assays that quantify concentrations of small molecules enable low-throughput targeted measurements, but are not suitable for CFE characterization. Importantly, multi-omics techniques can provide continuous, untargeted measurements of system composition beyond transcription and translation. While these techniques have already been used to study CFE systems (*13, 69–74*), multi-omics methods likely cannot complement routine characterization. Future characterization efforts should include easy-to-run, system-specific assays tailored for CFE systems.

Our work presents an important steppingstone to better characterization and understanding of CFE systems. We do not discourage spectrophotometric measurements of sfGFP at a single time point as a method of quality control. Instead, we aim to complement this measurement with our measurement framework. Further, we believe this framework can and should complement application-specific quality control. With our measurement framework and continuously evolving efforts, we believe we can realize the full potential of CFE systems.

## Materials and Methods

**Supplementary File 1** describes the preparation of nucleic acids, extracts, and CFE reactions in detail, and includes catalog numbers for all CFE reagents and materials (**Table S1**).

**Supplementary File 2** includes annotated sequences of all DNA sequences used in this work, and values computed for all reaction metrics.

### Materials

- All bacteria growth media reagents were purchased from Millipore Sigma.
- LB medium (10 g/L sodium chloride, 5 g/L yeast extract, and 10 g/L tryptone) was used to grow bacteria for plasmid cloning and extraction.
- 2xYTP medium (5 g/L sodium chloride, 10 g/L yeast extract, 16 g/L tryptone, 40 mmol/L potassium phosphate dibasic, and 22 mmol/L potassium phosphate monobasic) was used to grow bacteria for extract preparation.
- Kanamycin (50 μg/mL) was used for antibiotic selection.
- T4 polynucleotide kinase, T4 DNA ligase, T5 exonuclease, Phusion High-Fidelity PCR Master Mix, Gibson Assembly Master Mix, and DpnI—all purchased from New England Biolabs—were used for plasmid cloning.
- E.Z.N.A. Plasmid DNA Mini Kit (Omega Bio-tek) was used to extract plasmids for sequence verification.
- E.Z.N.A. Plasmid DNA Midi Kit (Omega Bio-tek) was used to extract plasmids for use in CFE reactions.
- T7 RNA Polymerase (Thermo Scientific) was used in *in vitro* transcription.
- RNA Clean & Concentrator-25 Kit (Zymo Research) was used to purify and concentrate mRNAs generated via *in vitro* transcription.
- E-Gel™ 1 % Agarose Gels with SYBR™ stain(Invitrogen) were used to visualize DNA.
- E-Gel™ 2 % Agarose Gels with EX stain (Invitrogen) were used to visualize mRNA.
- Quick Start™ Bradford Protein Assay Kit 1 (Bio-rad) and Nunc™ MicroWell™ 96-Well Microplates (Thermo Fisher) were used to run Bradford assays.
- The NIST-traceable fluorescein standard (Invitrogen) was used to generate a calibration curve for sfGFP fluorescence measurements. The fluorescein was dissolved in a sodium borate (Sigma-Aldrich) buffer.
- Atto 590 dye (Millipore Sigma) was used to generate a calibration curve for Pepper-HBC620 fluorescence measurements.
- Clear 384-well plates with clear, flat bottoms (Greiner Bio-one, 784101) were used to run CFE reactions.
- Black 384-well plates with clear, flat bottoms (Corning, 3544) were used in the experiment described in Figure 6.
- 384-well plates were covered with a clear adhesive film (Fisher Scientific, 08408240) during incubation at 37 °C.
- Commercial CFE systems (NEBExpress and PURExpress) were purchased from New England Biolabs.
- Sonicators Q125 and Q700 (Qsonica) were used to lyse cells for extract preparation. The Q700 sonicator was fitted with a 24-tip horn (Qsonica, 4579).
- All CFE reagents are listed in **Table S4**.

### Bacterial strains

*Escherichia coli* K12 DH10B was used for plasmid assembly and extraction. BL21 Star (DE3) and BL21 (DE3) were used for extract preparation.

### Plasmids and plasmid assembly

Plasmid pJL1 was a gift from Michael Jewett (Addgene plasmid #69496). All other plasmids used in this study were generated via either Gibson assembly (New England Biolabs) or inverse PCR. To assemble plasmids via Gibson assembly, insert and backbone fragments were amplified via PCR and purified. 20 µL Gibson reactions were conducted with a 3:1 molar ratio of insert to backbone PCR, and (4 to 5) ng of backbone PCR per µL of reaction. Reactions were incubated at 50 °C for 1 h, and 3.5 µL of the reaction were then used to transform chemically competent DH10B cells. Transformants were screened via colony PCR. To assemble plasmids via inverse PCR, DNA primers were phosphorylated with T4 polynucleotide kinase following the manufacturer’s protocol. Phosphorylated primers were then used to amplify the plasmid backbone via PCR. The PCR product was purified, and 100 ng of the PCR were subsequently used in a 20 µL ligation reaction with T4 DNA ligase incubated at room temperature for 2 h. 3 µL of the ligation product were used to transform chemically competent DH10B cells. Transformants were screened via colony PCR. Gene fragments encoding Pepper aptamers were purchased from Integrated DNA Technologies (IDT) as gBlocks (**Supplementary File 2**) and subsequently used to insert each aptamer into pJL1 via Gibson assembly. Pepper aptamer sequences were taken from Mumbleau *et al*.(*21*), but included two point mutations in one of the Pepper units to reduce sequence complexity and increase the likelihood of successful synthesis (**Supplementary File 2**). The P_J23100_ promoter was obtained from the Anderson promoter collection in the Standard Registry of Biological Parts.

### DNA Sequencing

For preliminary cloning verification, relevant regions of purified plasmid DNA underwent Sanger Sequencing by Psomagen. Prior to use in CFE reactions, all plasmids underwent whole-plasmid sequencing by Plasmidsaurus using Oxford Nanopore Technology with custom analysis and annotation.

### Extract Preparation

The extract preparation protocol was adapted from Sun *et al*.(*6*) and Kwon *et al.*(*25*) and divided into a 4 d protocol.

**Day 1**: An *E. coli* BL21 Star (DE3) glycerol stock was streaked onto an LB plate and incubated at 37 °C for 16 h.

**Day 2**: 50 mL of LB were inoculated with one *E. coli* BL21 Star (DE3) colony at 37 °C and 250 rpm for 16 h in a baffled 250 mL flask.

**Day 3**: 400 mL of 2xYTP were inoculated with 25 mL of the overnight culture at 37 °C, 250 rpm. About 1.5 h into growth, 0.4 mmol/L isopropyl β-d-1-thiogalactopyranoside (IPTG) was added to the culture to induce expression of T7 RNA polymerase. Upon reaching the target OD600 of 1.6 to 1.7, cells were harvested by centrifugation at 4 °C, 2700 xg, for 15 min, and washed three times with S30 buffer (10 mmol/L Tris acetate, 14 mmol/L magnesium acetate, and 60 mmol/L potassium acetate, pH adjusted to 8.2 with 5 mol/L potassium hydroxide, with 2 mmol/L dithiothreitol added immediately before use), with a centrifugation step between washes. After the last wash step, the mass of the cell pellet was determined, and cells were stored at - 80 °C.

**Day 4:** Cells were thawed on ice, then resuspended in 1 mL of S30 buffer per g of cells. The cellular resuspension was divided into 1 mL aliquots in 1.5 mL microcentrifuge tubes. The cellular resuspension was then lysed on ice using a Q125 sonicator (Qsonica) with a 3.175 mm diameter probe, 20 kHz frequency, 50 % amplitude, and cycles of 10 s on and 10 s off, delivering (260 ± 10) J after 5 cycles. At the start of each cycle, the tip of the probe was positioned close to the bottom of the tube; approximately 5 times per cycle, the tube was moved down slowly such that the tip reached the 0.5 mL mark of the tube and then reverted to its original position. Immediately after lysis, 3 mmol/L DTT were added to each tube. Lysed cells were centrifuged at 4 °C, 12000 xg, for 15 min. The supernatant was consolidated into a single tube, mixed by inversion, then divided into 2 mL aliquots in 14 mL culture tubes and allowed to shake at 37 °C, 250 rpm, for 80 min—this is the runoff reaction. The sample was then centrifuged at 4 °C, 12000 xg, for 15 min. The supernatant (*i.e.*, the final extract to be used in CFE reactions) was consolidated into one tube, mixed by inversion, divided into 200 µL aliquots, and stored at - 80 °C for future use.

To demonstrate reproducibility between extract batches, we measured transcription and translation dynamics in EXT4, an extract nominally identical to EXT3 but prepared on a different day (**Figure S13**). At most concentrations of pFP34 DNA tested, the differences in sfGFP and Pepper signals between the two extracts were smaller than the differences among extracts prepared under different conditions, indicating that the differences measured among extracts prepared in house (**Figure 4**) arose primarily from biological differences and not batch-to-batch variability.

### Cell-Free Reactions

Cell-free reactions in extracts were prepared as previously described(*31*). Unless otherwise specified, reactions contained 10 mmol/L magnesium glutamate, 10 mmol/L ammonium glutamate, 133 mmol/L potassium glutamate, 1.2 mmol/L ATP, 0.85 mmol/L GTP, 0.85 mmol/L CTP, 0.85 mmol/L UTP, 0.034 mg/mL folinic acid, 0.171 mg/mL tRNA from *E. coli* MRE 600, 0.33 mmol/L nicotinamide adenine dinucleotide, 0.2667 mmol/L coenzyme A, 4 mmol/L sodium oxalate, 1 mmol/L putrescine, 1.5 mmol/L spermidine, 50 mmol/L HEPES, 2 mmol/L each of the twenty standard amino acids, 0.03 M PEP, 27 % extract by volume, 5 µmol/L HBC620 dye, and a nucleic acid template concentration specified for each experiment (**Table S1**).

**Section III** of **Supplementary File 1** describes in detail how reactions were assembled. Unless otherwise specified, reactions were run in 3 10 µL technical replicates in clear 384-well plates in a BioTek Synergy Neo2 plate reader at 37 °C. Measurements of sfGFP (excitation: 485 nm, emission: 510 nm, bandwidth: 10 nm, gain: 75 or 50) and the Pepper-HBC620 complex (excitation: 577 nm, emission: 620 nm, bandwidth: 20, gain: 100) were taken every 4 min from the bottom of the plate.

### In vitro transcription

Prior to *in vitro* transcription, plasmid DNA was linearized to remove the origin of replication and the antibiotic resistance gene (**Table S2**). *In vitro* transcription reactions to generate mRNAs contained 1 µg of linear DNA template, 2 mmol/L each of ATP, GTP, CTP and UTP, 0.6 U/µL T7 RNA polymerase (Thermo Scientific), and the appropriate transcription buffer provided by the manufacturer. Reactions were run in 100 µL volumes at 37 °C for 2 h in a thermocycler. Each reaction was then treated with 0.1 U/µL DNase I for 15 min at room temperature prior to purification and elution in 30 µL nuclease-free water.

### Bradford Assay

The assay was run following the manufacturer’s protocol. In short, a calibration curve of bovine serum albumin (BSA) at (0, 1.25, 5, 7.5, 10) µg/mL was prepared by adding BSA to wells of a clear 96-well plate containing Bradford reagent. Bacterial extracts were mixed with Bradford reagent to a final dilution factor of 3000. The plate was incubated at room temperature for 10 min, then placed inside a BioTek Synergy Neo2 plate reader to enable measurement of absorbance at 595 nm. Extract absorbances were compared to the BSA calibration curve to determine the total protein concentration.

### Calibration curves

#### NIST-traceable fluorescein standard (NFS)

A 0.1 mol/L stock of sodium borate buffer (pH adjusted to 9.5 with 1 mol/L NaOH) was prepared. 50 µmol/L NFS was diluted to (15, 12.5, 10, 7.5, 5, 3.75, 1.875, 0.9375, 0.4688, 0.2344, 0.1172, 0.0586, 0.0293) µmol/L in the sodium borate buffer. For each CFE reaction format, fluorescein dilutions were added to wells of the appropriate 384-well plate at the same volume as the CFE reaction, and fluorescence was measured at the settings used for sfGFP.

#### Atto 590

Solutions of Atto 590 were prepared at (25, 20, 17.5, 15, 10, 7.5, 5, 2.5) µmol/L in 100 % DMSO. For each CFE reaction format, fluorescein dilutions were added to wells of the appropriate 384-well plate at the same volume as the CFE reaction, and fluorescence was measured at the settings used for the Pepper-HBC620 complex.

Linear regression was implemented for each fluorescein using the “linregress” function from Python’s SciPy library. To convert fluorescence measurements from arbitrary units to Molecules of Equivalent Soluble Fluorochrome (MESF) in µmol/L, a line with the slope of the linear regression and a y-intercept corresponding to the fluorescence value generated by a sample lacking any fluorescein (*i.e.,* containing solvent only) was used. The y-intercept of the linear regression was not used because it was physically unrealistic: it was negative for NFS, and orders of magnitude greater than 0 for Atto 590. **Figure S1** shows calibration curves for the two fluorochromes in both microplates used in this study.

### Data analysis and visualization

**Supplementary File 1, Section IV** describes data analysis in detail. In summary, all experimental data were processed in the Jupyter Notebook interface using Python scripts and leveraging the Pandas and Bokeh libraries. The fluorescence intensity values of three technical replicates were averaged. The averages and their corresponding standard deviations were then converted to Molecules of Equivalent Soluble Fluorochrome (MESF), and the average MESF value of a reaction lacking exogenous nucleic acid templates was subtracted from the average MESF value of all other samples in the experiment. Plots were generated using the Bokeh data visualization library, and final Figures were made in Adobe Illustrator 2023.

## Author Contributions

**FP:** Conceptualization, Methodology, Investigation, Data curation, Formal analysis, Software, Visualization, Writing – original draft

**CS:** Methodology, Writing – review and editing

**EAS:** Conceptualization, Methodology, Supervision, Writing – review and editing

**ER:** Conceptualization, Methodology, Supervision, Writing – review and editing

## Acknowledgements

We acknowledge the financial support of the National Institute of Standards and Technology. The authors thank Samuel Schaffter and Tyler Goshia for helpful comments during the drafting process, and Zoila Jurado for her valuable help with data processing and visualization. The authors also thank Michael Jewett for plasmid pJL1.

## Disclaimer

Certain commercial entities, equipment, or materials may be identified in this document to describe an experimental procedure or concept adequately. Such identification is not intended to imply recommendation or endorsement by the National Institute of Standards and Technology, nor is it intended to imply that the entities, materials, or equipment are necessarily the best available for the purpose. Official contribution of the National Institute of Standards and Technology; not subject to copyright in the United States.

## Supplementary Information

*Supplementary File 1* (.docx): Additional details on nucleic acid templates used in each experiment, measurement calibration, extract preparation, and reaction formulation and assembly; additional data.

*Supplementary File 2* (.xlsx): Annotated sequences for all plasmids, Pepper aptamer scaffolds, and gBlocks used in this study; computed reaction metrics for all experiments.

