## Supplementary File 1 for "Characterizing Cell-Free Transcription and Translation Dynamics with Nucleic Acid-Based Assays"

**Supplementary File I**

### I. Nucleic acid templates

**Table S1: Nucleic acid templates used in each experiment.**

| **Figure** | **Nucleic acid template** | **Type** | **Concentration (nmol/L)** |
| --- | --- | --- | --- |
| 2A-C, left | pJL1,  pFP34, pFP35 | DNA | 5 |
| 2A-C, center | pFP34, pFP34∆sfGFP, pFP34∆RBS | DNA | 5 |
| 2A-C, right | pFP35, pFP35∆sfGFP, pFP345∆RBS | DNA | 5 |
| 3A-C, left | pJL1, pFP34, pFP35 | RNA | 300 |
| 3A-C, center | pFP34, pFP34∆sfGFP, pFP34∆RBS | RNA | 300 |
| 3A-C, right | pFP35, pFP35∆sfGFP, pFP345∆RBS | RNA | 300 |
| 4B-D, left | pFP34 | DNA | 5 |
| 4B-D, center | pFP34 | DNA | 5 |
| 4B-D, right | pFP35 | DNA | 5 |
| 5B | pFP24 | DNA | 5 |
| 5C | pFP59 | DNA | 5 |
| 5D | pFP60 | DNA | 5 |
| 6A-B, left | pFP34 | DNA | 5 |
| 6A-B, center | pFP34 | DNA | 5 |
| 6A-B, right | pFP35 | DNA | 5 |
| 7A | pFP34 | DNA | 5 |
| 7B | pFP35 | DNA | 5 |

**Table S2: Primers used to linearize plasmids prior to *in vitro* transcription.**

|  | **Primer sequence (5′ to 3′)** |
| --- | --- |
| Forward primer | TGTCGGGTTTCGCCACCTC |
| Reverse primer | CAGTTTCATTTGATGCTCGATGAGTTTTTC |

### II. Measurement calibration

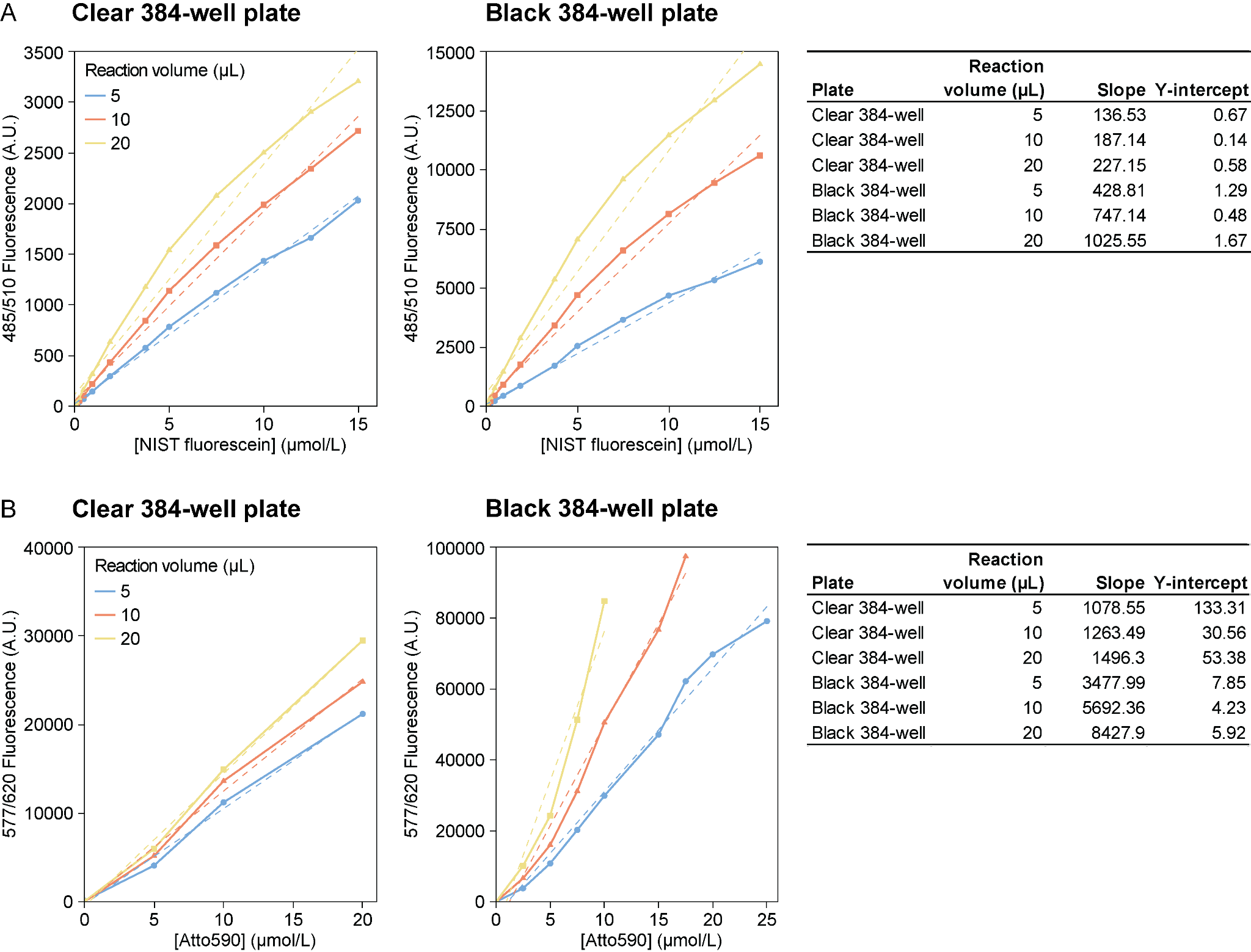

**Figure S1.** Calibration curves for the fluorochromes used in this work. **(A)** NIST-traceable fluorescein standard and **(B)** Atto 590, each at 5, 10 or 20 µL reaction volumes in either a clear 384-well plate (Greiner Bio-One, 784101) or a black 384-well plate (Corning, 3544). The dashed lines were generated via linear regression. The tables indicate the slope and y-intercept values used to convert fluorescence intensity values to Molecules of Equivalent Soluble Fluorochrome (MESF) in µmol/L in all the experiments presented in this study. The slope value was taken directly from the linear regression, and the fluorescence value for a measurement of buffer (sodium borate of DMSO) lacking the fluorochrome was used as the y-intercept.

### III. Cell-free extracts and reactions

*Bacterial genotypes*

BL21 (DE3): F– *omp*T *hsdS_B_* (r_B_–, m_B_–) *gal dcm* (DE3)

BL21 Star (DE3): F–ompT hsdSB (rB–, mB–) gal dcm rne131 (DE3)

Where (DE3) = λ sBamHIo ∆EcoRI-B int::(lacI::PlacUV5::T7 gene1) i21 ∆nin5

*Extract preparation*

The Methods section of the manuscript describes the protocol used to prepare EXT3. To prepare the other extracts characterized in this study, we modified this baseline protocol, as listed in Table 1. Below are more details on the protocol modifications for each extract.

- **EXT10:** Prior to lysis, the cell resuspension was divided into 1 mL aliquots in 1.5 mL microcentrifuge tubes. Tubes were placed in a pre-cooled rack (Qsonica, 514), and samples were lysed simultaneously using a Q700 sonicator (Qsonica) with a 24-tip horn. Each tip has a 3.175 mm diameter, matching the single tip of the Q125 sonicator (Qsonica) used to prepare all other extracts. While Q700 was programmed with the same settings as Q125 (20 kHz frequency, 50 % amplitude, cycles of 10 s on and 10 s off), Q700 did not display the sonication energy input delivered by each tip; instead, it displayed the total energy input, which was lower than the target input per tube or tip (around 260 J) when divided by the number of tubes. Because tubes were placed in a rack during sonication, they could not be moved up and down, likely resulting in more uneven lysis than for other extracts. In addition, fewer than 24 tubes were used at a time, so tubes filled with water were placed in the empty slots in the rack. We previously noticed that samples on the edges of the rack had different lysis efficiencies than samples in interior slots of the rack; when preparing EXT10, we did not place samples on the edges of the rack to avoid this issue.
- **EXT12:** The extract was prepared from *E. coli* BL21 (DE3) cells lacking the Star (rne131) mutation.
- **EXT14:** After harvest, cells were resuspended and washed in S30A buffer with a pH adjusted to 10 instead of 7.7. 1 mL of this buffer was added per g of cells prior to lysis.
- **EXT15:** Cell cultures were allowed to grow until reaching an optical density of 2.0 at a wavelength of 600 nm (OD600).
- **EXT16:** Cell cultures were allowed to grow until reaching an OD600 of 2.2.
- **EXT17:** After harvest, cells were resuspended and washed in S30A buffer instead of S30 buffer. 1 mL of S30A was added per g of cells prior to lysis.
- **EXT18:** After harvest, cells were resuspended and washed in S30A buffer instead of S30 buffer. 1 mL of S30A was added per g of cells prior to lysis. After cell lysis and centrifugation, the supernatant was isolated and used directly in cell-free reactions without undergoing a runoff reaction.
- **EXT19:** Cell cultures were allowed to grow for 24 h prior to harvest.
- **EXT20:** After cell lysis and centrifugation, the supernatant was isolated and used directly in cell-free reactions without undergoing a runoff reaction.
- **EXT23:** During cell growth, no IPTG was added to induce expression of T7 RNAP.

*Reaction formulation*

To facilitate reaction assembly and ensure reagent integrity, reagents for cell-free reactions were prepared by grouping them into multiple concentrated solutions. A stock solution of HBC620 was prepared by dissolving HBC620 powder in DMSO.

**Table S3. Cell-free reaction formulation.**

| **Reagent** | **Composition** | **Stock concentration** | **Final concentration** | **Unit** |
| --- | --- | --- | --- | --- |
| **Solution I** | Magnesium glutamate | 150 | 10 | mmol/L |
|  | Potassium glutamate | 2000 | 133.3 | mmol/L |
|  | Ammonium glutamate | 150 | 10 | mmol/L |
| **Solution II** | ATP | 18 | 1.2 | mmol/L |
|  | CTP | 12.75 | 0.85 | mmol/L |
|  | GTP | 12.75 | 0.85 | mmol/L |
|  | UTP | 12.75 | 0.85 | mmol/L |
|  | Folinic acid | 0.51 | 0.034 | mg/mL |
|  | tRNA | 2.565 | 0.171 | mg/mL |
| **Solution III** | NAD | 3.96 | 0.33 | mmol/L |
|  | Coenzyme A | 3.204 | 0.267 | mmol/L |
|  | Sodium oxalate | 48 | 4 | mmol/L |
|  | Putrescine | 12 | 1 | mmol/L |
|  | Spermidine | 18 | 1.5 | mmol/L |
|  | HEPES | 600 | 50 | mmol/L |
| **Amino acids** | 20 standard AAs | 50 | 2 | mmol/L |
| **Energy molecule** | PEP | 1000 | 30 | mmol/L |
| **HBC620** | HBC620 | 1000 | 5 | µmol/L |
| **Extract** | Extract | - | 26.7 | % (v/v) |

In the experiment described in Figure 7, pyruvate was added to the relevant reactions at a concentration of 30 mmol/L in addition to or in place of PEP.

*Reaction preparation*

Nucleic acid templates and water were mixed in PCR tubes on ice. A “master mix” of solutions I-III, amino acids, HBC620 dye, and the energy molecule was prepared separately in a microcentrifuge tube, mixed by vortexing, then divided into PCR tubes. Reactions were mixed by vortexing, then 10 µL of each reaction were added to each of 3 wells (constituting technical triplicates) of a 384-well plate stored at room temperature.

*Reaction formats*

Unless otherwise specified, reactions were run at 10 µL volumes in a 384-well plate incubated at 37 ˚C in a microplate reader without shaking. Plates were sealed with an adhesive film. Plate dimensions and drawings can be found on the manufacturer’s website. Two types of plate were used:

- Greiner Bio-one, catalog number 784101: non-binding, clear plate with a clear, flat bottom, and a working volume of 4 µL to 25 µL. Each well can hold up to 28 µL.
- Corning, catalog number 3544: non-binding, black plate with a clear, flat bottom, and a working volume of 5 µL to 40 µL. Each well can hold up to 50 µL.

**Table S4: Reagents used to prepare extracts and cell-free reactions.**

| **Reagent** | **Manufacturer** | **Catalog number** |
| --- | --- | --- |
| Tryptone | Fisher Scientific | BP1421-500 |
| Sodium chloride | Millipore Sigma | S3014-500G |
| Yeast extract | Fisher Scientific | BP1422-500 |
| Potassium phosphate dibasic | Millipore Sigma | 60353-250G |
| Potassium phosphate monobasic | Millipore Sigma | P9791-100G |
| Isopropyl β-D-1-thiogalactopyranoside (IPTG) | Millipore Sigma | I6758-5G |
| Trizma® base | Millipore Sigma | T6066-500G |
| Acetic acid | Millipore Sigma | 695092-100ML |
| Dithiothreitol (DTT) | Thermo Fisher (Invitrogen) | 15508013 |
| HBC620 | MedChemExpress | HY-133520 |
| Coenzyme A sodium salt hydrate (CoA) | Millipore Sigma | C3144-25MG |
| β-Nicotinamide adenine dinucleotide (NAD) | Millipore Sigma | N8535-15VL |
| tRNA from *E. coli* MRE600 | Millipore Sigma (Roche) | 10109541001¹ |
| L-Glutamic acid hemimagnesium salt tetrahydrate | Millipore Sigma | 49605-250G |
| L-Glutamic acid potassium salt monohydrate | Millipore Sigma | G1149-500G |
| L-Glutamic acid monoammonium salt | Toronto Research Chemicals | TRC-G597140 |
| Folinic acid | Millipore Sigma | F7878-100MG |
| Spermidine | Millipore Sigma | 85558-5G |
| 1,4-Diaminobutane (putrescine) | Millipore Sigma | D13208-25G |
| HEPES | Thermo Fisher | 11344041 |
| Potassium oxalate monohydrate (oxalic acid) | Millipore Sigma | P0963-100G |
| NTP Set | ThermoFisher | R0481 |
| Potassium hydroxide | Fisher Scientific | SP236-500 |
| Phosphoenolpyruvate (PEP) | Millipore Sigma | 860077-250MG |
| Pyruvate | Millipore Sigma | P5280-25G |
| L-Aspartic Acid | Millipore Sigma | A7219-100G |
| L-Valine | Millipore Sigma | V0500-25G |
| L-Tryptophan | Millipore Sigma | T0254-25G |
| L-Phenylalanine | Millipore Sigma | P2126-100G |
| L-Isoleucine | Millipore Sigma | I2752-25G |
| L-Leucine | Millipore Sigma | L8000-25G |
| L-Cysteine | Millipore Sigma | C7352-25G |
| L-Methionine | Millipore Sigma | M9625-25G |
| L-Alanine | Millipore Sigma | A7627-100G |
| L-Arginine | Millipore Sigma | A8094-25G |
| L-Asparagine | Millipore Sigma | A0884-25G |
| Glycine | Millipore Sigma | G7126-100G |
| L-Glutamine | Millipore Sigma | G3126-250G |
| L-Histidine | Millipore Sigma | H8000-25G |
| L-Lysine | Millipore Sigma | L5501-25G |
| L-Proline | Millipore Sigma | P0380-10G |
| L-Serine | Millipore Sigma | S4500-100G |
| L-Threonine | Millipore Sigma | T8625-25G |
| L-Tyrosine | Millipore Sigma | T3754-100G |

^1^tRNA from *E. coli* MRE600, previously manufactured by Roche and widely used for cell-free gene expression, has been discontinued. Sigma Aldrich provided another commercially available tRNA product, but from *E. coli* W (catalog number R1753). We have observed similar sfGFP yields from pJL1 with the two tRNA sources, although stronger cell-free expression is reported with Roche’s tRNA product in a reconstituted system prepared in house*(1)*. At the time this manuscript was written, Sigma Aldrich’s tRNA product was no longer available for purchase.

### IV. Data processing and analysis

*Data processing*

We opted to use “signal” to refer to sfGFP and Pepper expression data because the measurable fluorescence intensity, which we later converted to Molecules of Equivalent Soluble Fluorochrome (MESF), encompasses multiple processes. For sfGFP, the increase in fluorescence intensity reflects the transcription, translation, and maturation of sfGFP, including protein folding and formation of the fluorescent chromophore. For the Pepper RNA aptamer, the increase in fluorescence intensity results from transcription, folding, and binding to the HBC620 dye.

We used a Python script to process sfGFP and Pepper-HBC620 fluorescence intensity data and compute metrics for cell-free reactions. Our Python script followed the steps summarized below.

1) Compute the average of three technical replicates and their standard deviation.
2) Convert the average fluorescence intensity and corresponding standard deviation to MESF.
3) For each reaction, subtract the MESF for a reaction containing all reagents except added nucleic acid templates from the average MESF.
4) Compute reaction metrics.

*Computation of reaction metrics*

The values of all computed reaction metrics are listed in the Reaction Metrics sheet of Supplementary File 2. For sfGFP expression data, we computed the maximum signal, the time to the maximum signal, the maximum rate of signal increase, and time to the maximum rate of signal increase. Once the sfGFP signal plateaued, we did not observe a decrease in signal in any experiments. For Pepper expression data, we included two additional metrics—the maximum rate of signal decrease and the time to reach that rate—to account for the degradation of Pepper RNA.

To calculate reaction metrics, we first generated plots for the derivatives of the sfGFP and Pepper signals. We used the “signal” function from the SciPy library with a Savitzky-Golay filter (“savgol_filter”) to generate an array of derivative values. As the function’s inputs, we specified a window length of 5, a polynomial order of 1, a derivative order of 1, and a delta value equal to the time interval between measurements. We then computed the metrics below.

- **Maximum signal**: Computed as the maximum average MESF value, and also as the signal measured when two sequential derivative values differed by less than the associated standard deviation.
- **Time to maximum signal**: The time at which the maximum signal was measured.
- **Maximum rate of signal increase**: The maximum derivative value.
- **Time to the maximum rate of signal increase**: The time at which the maximum rate of signal increase was measured.
- **Maximum rate of signal decrease:** The minimum derivative value.
- **Time to the maximum rate of signal increase**: The time at which the maximum rate of signal decrease was measured.

For sfGFP measurements, the maximum signal is often referred to as the “endpoint” signal, due to negligible protein degradation in BL21 extracts and PURE systems. For Pepper measurements, the maximum signal marks the transition from a regime of signal increase to a regime of signal decrease, when RNA degradation exceeds Pepper aptamer transcription, folding, and binding to the HBC620 dye.

Measurements of both maximum signal and rate are needed to provide a complete picture of a reaction and enable accurate comparison of different systems. For example, reactions may have the same maximum rate of sfGFP signal increase, but one may sustain higher rates for longer than the other, resulting in a higher maximum sfGFP signal (Figure 4D, left). While the maximum rates of signal increase and decrease do not capture full reaction dynamics over the course of a reaction, these metrics simplified our analysis and provided additional insight into reaction longevity and tradeoffs between transcription and translation.

As we did for all other reactions, we computed the maximum rate of signal increase for reactions with RNA templates in EXT3, although the computed rate does not reflect an actual increase in signal. This value results from minor noise in the data.

*Temperature-related effects on fluorescence measurements*

In some experiments, measurements of sfGFP and Pepper signal exhibited a decrease in signal in the first (30 to 40) min of the reaction. Such a decrease is typical of the reaction format used in this work and can likely be attributed to the change in temperature as the reaction vessel equilibrates to 37 ˚C in the plate reader. This feature is more commonly observed for translation, which has a longer lag than transcription. However, we also observed an initial decrease in Pepper signal with most ∆sfGFP and ∆RBS DNA templates, likely because processes leading to an increase in Pepper signal (transcription, folding, and binding to the HBC620 dye) were not sufficiently strong to mask the temperature-related signal decrease (Figure 3A, center and right). This initial signal decrease and the subsequent signal increase interfered with the calculation of reaction metrics, so for certain reactions we excluded these data points from our analysis.

### V. Additional data

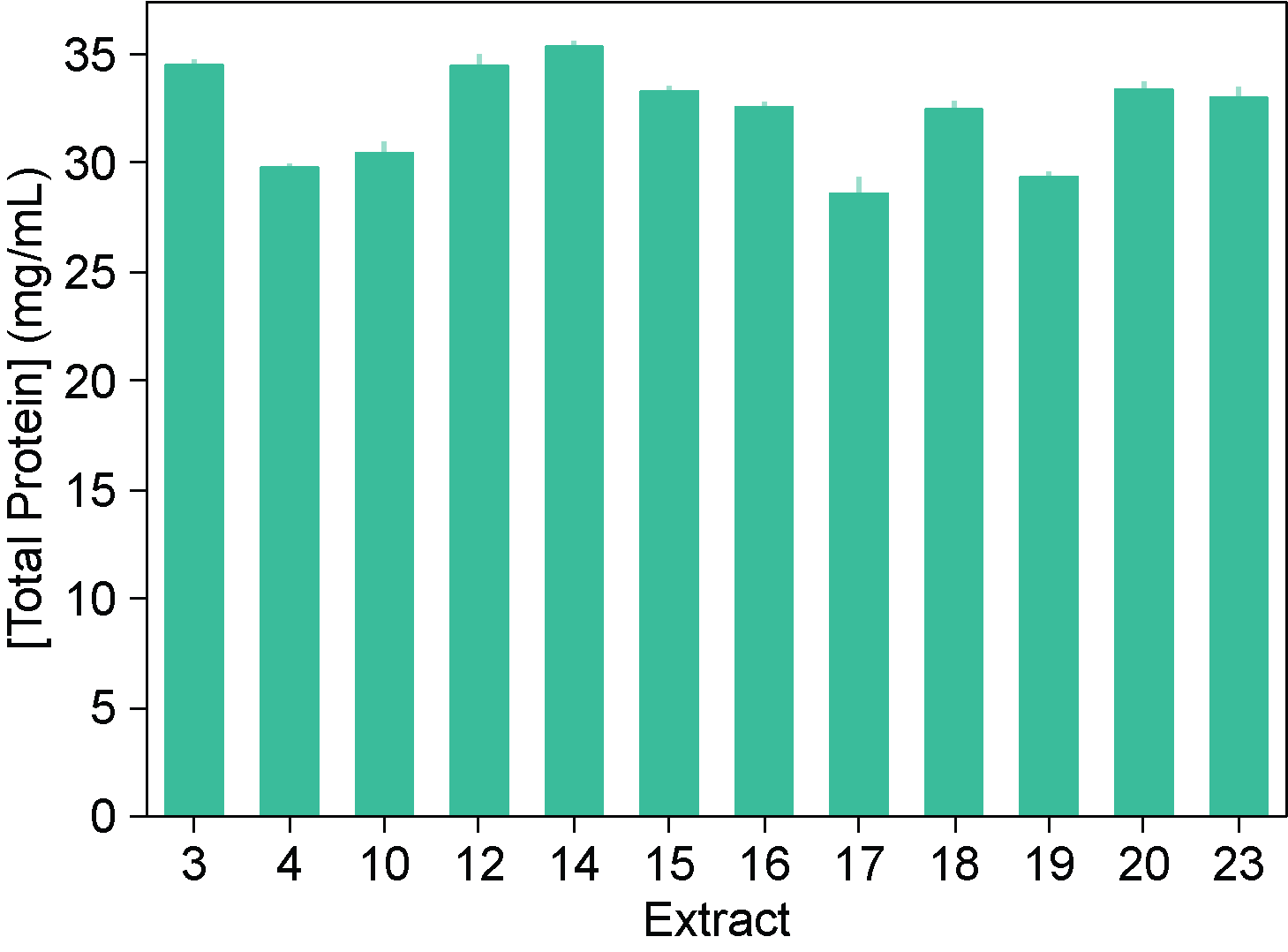

**Figure S2:** Total endogenous protein content of extracts prepared in house, measured via Bradford assay. The error bars indicate the standard deviation of three technical triplicates.

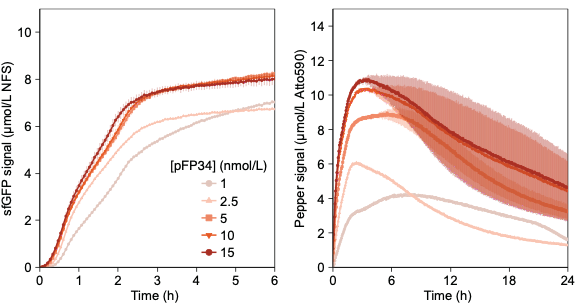

**Figure S3:** Measurements of translation and transcription dynamics at multiple concentrations of pFP34 and pFP35 DNA in EXT3. sfGFP measurements are reported in Molecules of Equivalent Soluble Fluorochrome (MESF) for a NIST-traceable fluorescein standard (NFS) and are shown for only the first 6 h of the reaction to highlight differences among reactions. Pepper mRNA measurements are reported in MESF for Atto 590. Note the difference in y-axis ranges. The error bars indicate the standard deviation of three technical triplicates.

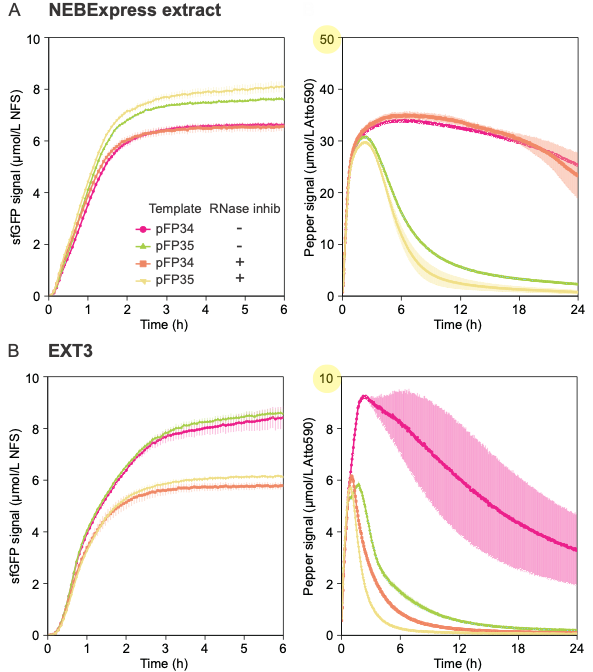

**Figure S4:** Measurements of transcription and translation dynamics in **(A)** NEBExpress extract and **(B)** EXT3 with and without added RNase inhibitor at 0.8 U/µL. All reactions include a DNA template at 5 nmol/L. sfGFP measurements are reported in Molecules of Equivalent Soluble Fluorochrome (MESF) for a NIST-traceable fluorescein standard (NFS) and are shown for only the first 6 h of the reaction to highlight differences among reactions. Pepper mRNA measurements are reported in MESF for Atto 590. Note the difference in y-axis ranges. The error bars indicate the standard deviation of three technical triplicates.

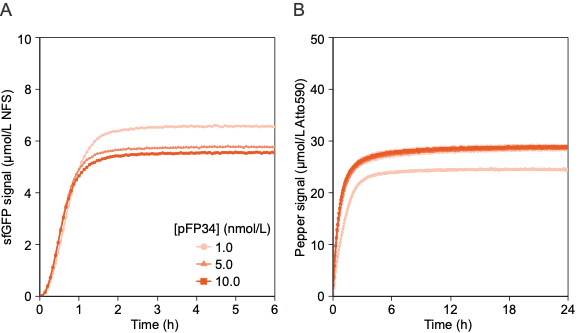

**Figure S5.** Measurements of **(A)** translation and **(B)** transcription dynamics in PURExpress from pFP34 DNA added at different concentrations. Although 1 nmol/L DNA yielded a higher sfGFP signal, 5 nmol/L DNA were used in the Figure 3 experiments to match the concentration used in extract-based reactions. sfGFP measurements are reported in Molecules of Equivalent Soluble Fluorochrome (MESF) of a NIST-traceable fluorescein standard (NFS) and are shown for only the first 6 h of the reaction, because the signal remains constant until 24 h. Pepper mRNA measurements are reported in MESF for Atto 590. The error bars indicate the standard deviation of three technical triplicates.

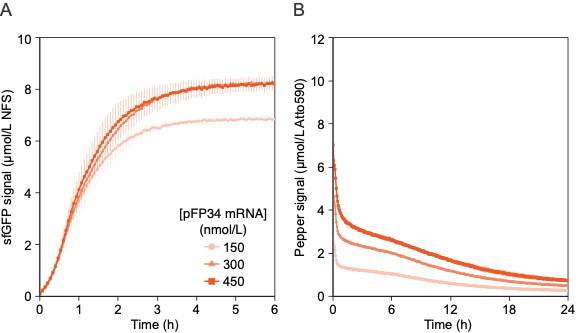

**Figure S6**. Measurements of **(A)** translation and **(B)** transcription dynamics in EXT3 from pFP34 mRNA added at different concentrations. A concentration of 300 nmol/L was used in other experiments with mRNA templates because it maximized sfGFP signal. sfGFP measurements are reported in Molecules of Equivalent Soluble Fluorochrome (MESF) of a NIST-traceable fluorescein standard (NFS) and are shown for only the first 6 h of the reaction, because the signal remains constant until 24 h. Pepper mRNA measurements are reported in MESF for Atto 590. The error bars indicate the standard deviation of three technical triplicates.

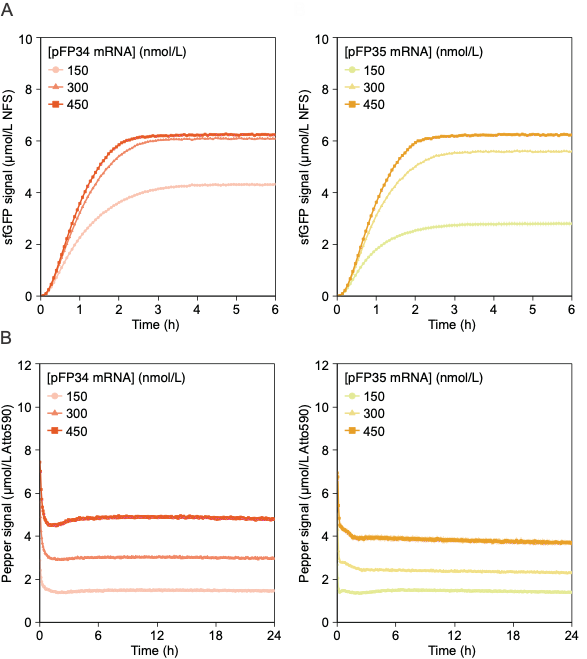

**Figure S7**. Measurements of **(A)** translation and **(B)** transcription dynamics in PURExpress from pFP34 and pFP35 mRNA templates added at different concentrations. A concentration of 300 nmol/L was used in other experiments with mRNA templates to match the concentration used in other CFE systems. sfGFP measurements are reported in Molecules of Equivalent Soluble Fluorochrome (MESF) of a NIST-traceable fluorescein standard (NFS) and are shown for only the first 6 h of the reaction, because the signal remains constant until 24 h. Pepper mRNA measurements are reported in MESF for Atto 590. The error bars indicate the standard deviation of three technical triplicates.

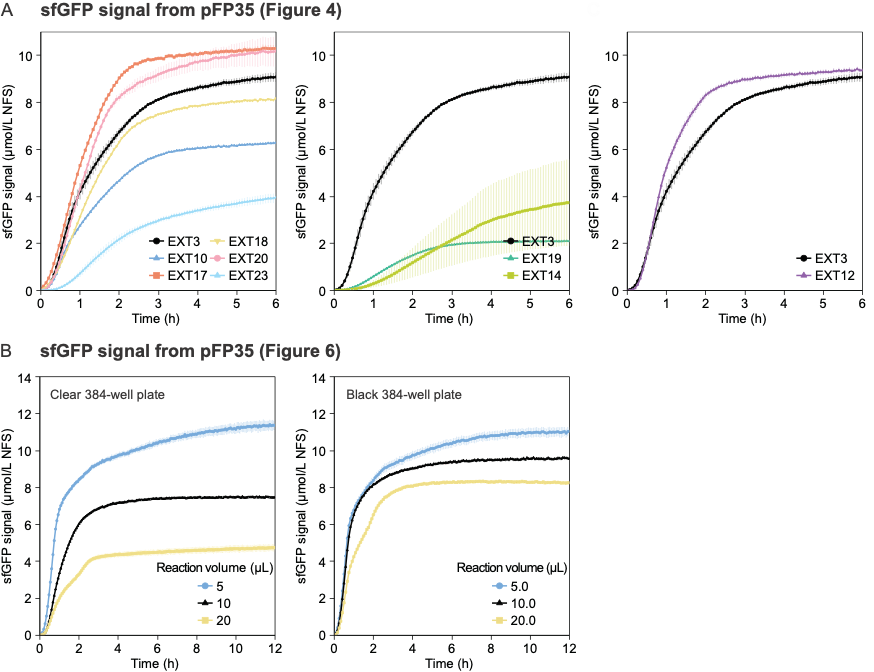

**Figure S8:** Measurements of translation dynamics with pFP35 as the DNA template corresponding to experiments shown in **(A)** Figure 4 and **(B)** Figure 6. These data were omitted from Figures 4 and 6 due to the similarity between the sfGFP signals generated with pFP34 and pFP35. sfGFP measurements are reported in Molecules of Equivalent Soluble Fluorochrome (MESF) of a NIST-traceable fluorescein standard (NFS). The error bars indicate the standard deviation of three technical triplicates.

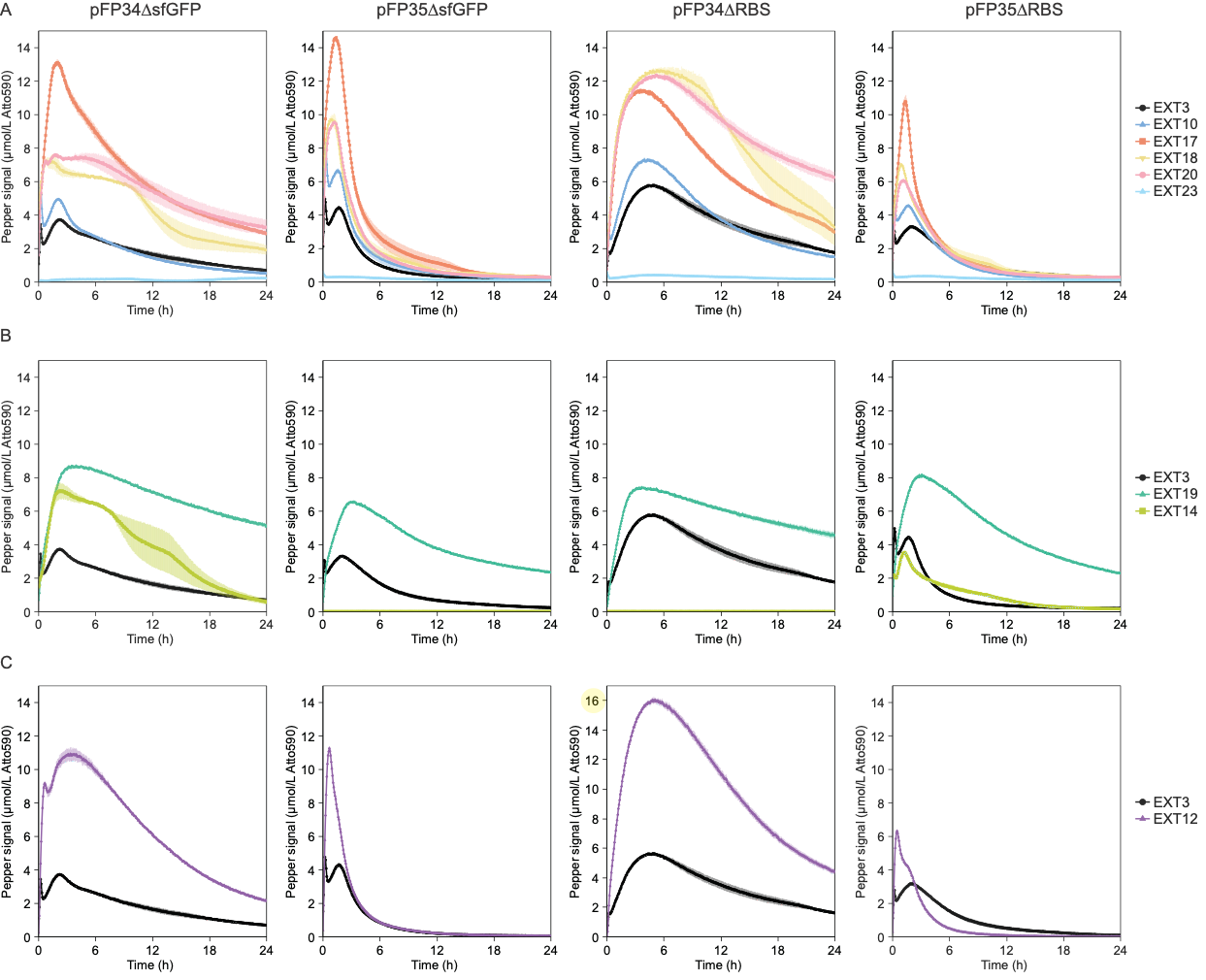

**Figure S9.** Measurements of transcription dynamics in extracts prepared in house **(B)** under different conditions, **(C)** under suboptimal conditions, and **(D)** from different *E. coli* host strains. Each panel includes the Pepper signal from pFP34∆sfGFP, pFP35∆sfGFP, pFP34∆RBS or pFP35∆RBS, with the relevant DNA template added at 5 nmol/L. Pepper mRNA measurements are reported in MESF for Atto 590. Note the different y-axis scale for pFP34∆RBS in (C). The error bars indicate the standard deviation of three technical triplicates.

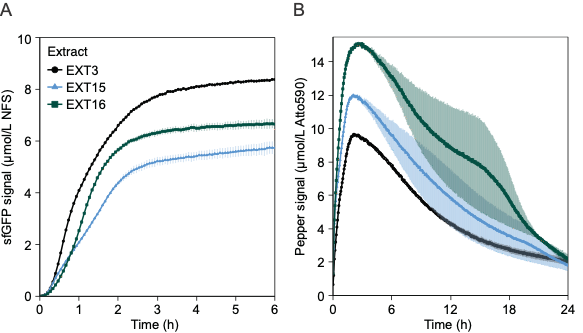

**Figure S10.** Measurements of **(A)** translation and **(B)** transcription dynamics in extracts derived from cells harvested in late-exponential phase (EXT15 and EXT16) with pFP34 as the DNA template. All reactions include pFP34 at 5 nmol/L. sfGFP measurements are reported in Molecules of Equivalent Soluble Fluorochrome (MESF) for a NIST-traceable fluorescein standard (NFS) and are shown for only the first 6 h of the reaction to highlight differences among extracts. Pepper mRNA measurements are reported in MESF for Atto 590. The error bars indicate the standard deviation of three technical triplicates.

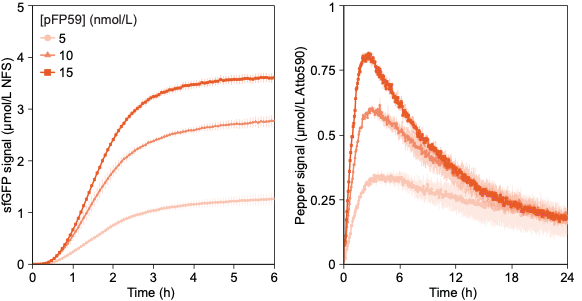

**Figure S11.** Measurements of translation (left) and transcription (right) dynamics in EXT3 at multiple concentrations of pFP59. sfGFP measurements are reported in Molecules of Equivalent Soluble Fluorochrome (MESF) for a NIST-traceable fluorescein standard (NFS) and are shown for only the first 6 h of the reaction to highlight differences among extracts. Pepper mRNA measurements are reported in MESF for Atto 590. The error bars indicate the standard deviation of three technical triplicates.

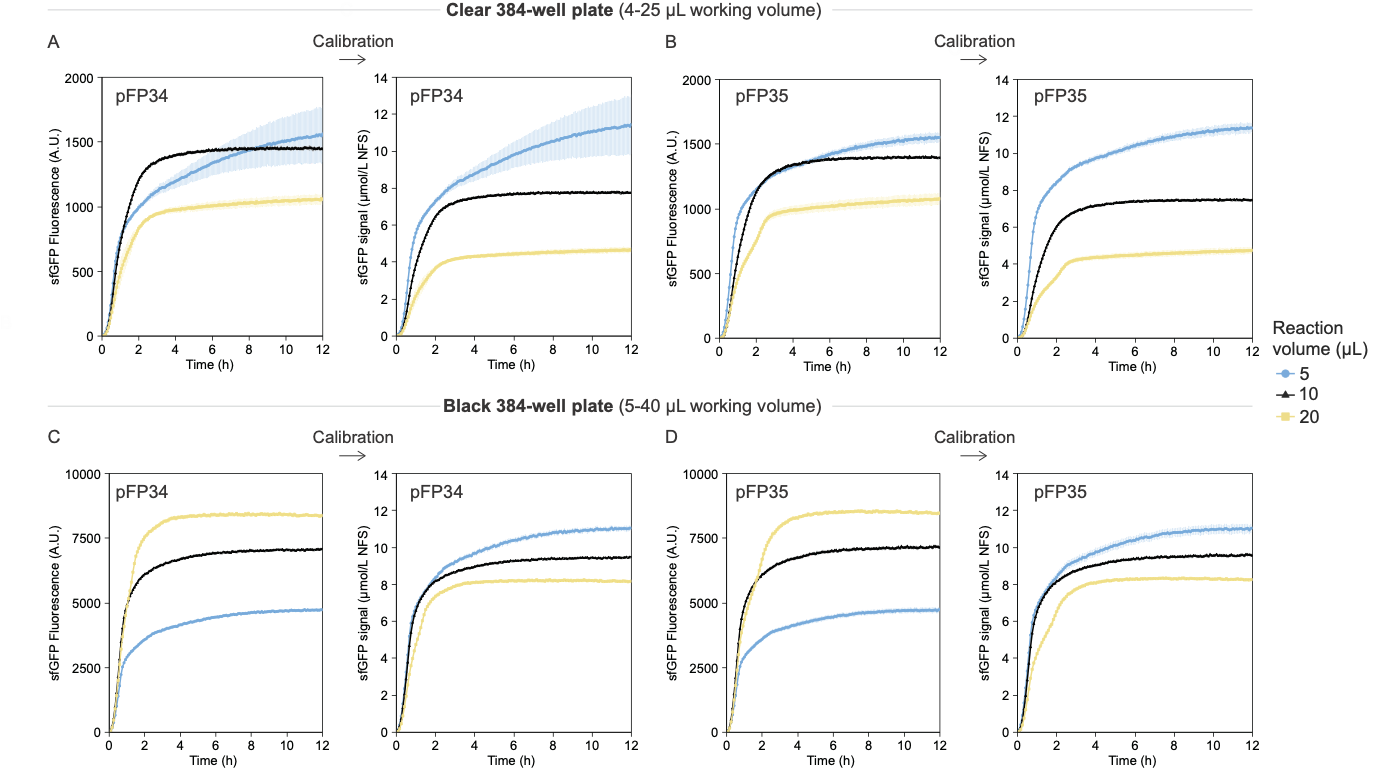

**Figure S12.** Measurements of sfGFP fluorescence intensity in arbitrary units (A.U.) generated in **(A,B)** a clear 384-well plate and **(C,D)** a black 384-well plate with pFP34 or pFP35 DNA templates at multiple reaction volumes. Fluorescence intensity values were converted to units of Molecules of Equivalent Soluble Fluorochrome (MESF) for a NIST-traceable fluorescein standard (NFS), and are displayed in Figure 6. Signal calibration changed the trend for sfGFP signal with increasing reaction volumes in the black 384-well plate. In all experiments, the DNA templates were added at 5 nmol/L. The error bars indicate the standard deviation of three technical triplicates.

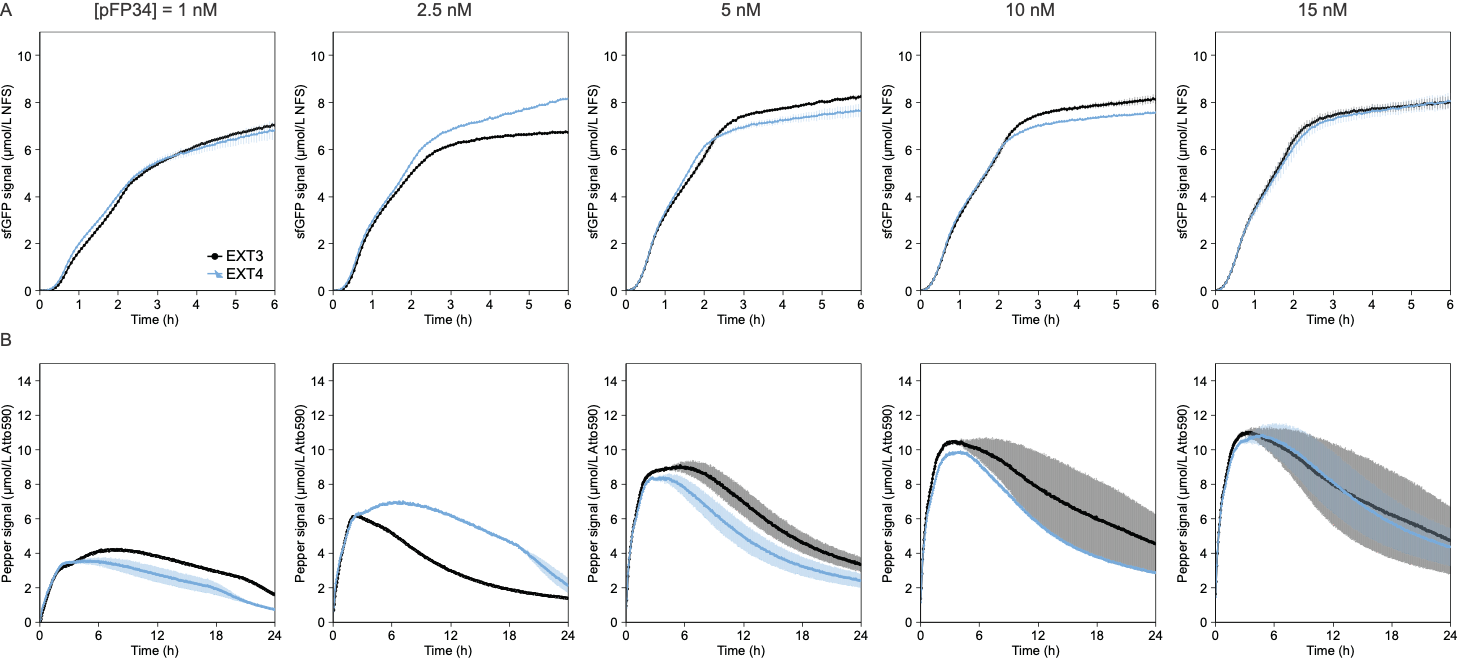

**Figure S13.** Measurements of **(A)** translation and **(B)** transcription dynamics from pFP34 DNA in EXT3 and EXT4, two nominally identical extracts prepared on different days. All reactions include pFP34 DNA at 5 nmol/L. sfGFP measurements are reported in Molecules of Equivalent Soluble Fluorochrome (MESF) for a NIST-traceable fluorescein standard (NFS) and are shown for only the first 6 h of the reaction. Pepper mRNA measurements are reported in MESF for Atto 590. The error bars indicate the standard deviation of three technical triplicates.

*Disclaimer*: Certain commercial entities, equipment, or materials may be identified in this document to describe an experimental procedure or concept adequately. Such identification is not intended to imply recommendation or endorsement by the National Institute of Standards and Technology, nor is it intended to imply that the entities, materials, or equipment are necessarily the best available for the purpose. Official contribution of the National Institute of Standards and Technology; not subject to copyright in the United States.
